# *Anopheles coluzzii* stearoyl-CoA desaturase is essential for adult female survival and reproduction upon blood feeding

**DOI:** 10.1101/2020.06.14.151019

**Authors:** Zannatul Ferdous, Silke Fuchs, Volker Behrends, Nikolaos Trasanidis, Dina Vlachou, George K. Christophides

## Abstract

Vitellogenesis and oocyte maturation require anautogenous female *Anopheles* mosquitoes to obtain a bloodmeal from a vertebrate host. The bloodmeal is rich in proteins that are readily broken down into amino acids in the midgut lumen and absorbed by the midgut epithelial cells where they are converted into lipids and then transported to other tissues including ovaries. The stearoyl-CoA desaturase (SCD) plays a pivotal role in this process by converting saturated (SFAs) to unsaturated (UFAs) fatty acids; the latter being essential for maintaining cell membrane fluidity amongst other housekeeping functions. Here, we report the functional and phenotypic characterization of SCD1 in the malaria vector mosquito *Anopheles coluzzii*. We show that RNA interference (RNAi) silencing of *SCD1* and administration of sterculic acid (SA), a small molecule inhibitor of SCD1, significantly impact on the survival and reproduction of female mosquitoes following blood feeding. Microscopic observations reveal that the mosquito thorax is quickly filled with blood, a phenomenon likely caused by the collapse of midgut epithelial cell membranes, and that epithelial cells are depleted of lipid droplets and oocytes fail to mature. Transcriptional profiling shows that genes involved in protein, lipid and carbohydrate metabolism and immunity-related genes are the most affected by *SCD1* knock down (KD) in blood-fed mosquitoes. Metabolic profiling reveals that these mosquitoes exhibit increased amounts of saturated fatty acids and TCA cycle intermediates, highlighting the biochemical framework by which the *SCD1* KD phenotype manifests as a result of a detrimental metabolic syndrome. Accumulation of SFAs is also the likely cause of the potent immune response observed in the absence of infection, which resembles an auto-inflammatory condition. These data provide insights into mosquito bloodmeal metabolism and lipid homeostasis and could inform efforts to develop novel interventions against mosquito-borne diseases.

## Introduction

In preparation for reproduction, anautogenous female mosquitoes must ingest vertebrate blood to improve their energy status and to stimulate vitellogenesis. This induces massive mobilization of lipids from the fat body to the ovaries where they serve as the principle energy source for the maturing oocytes and the developing embryos [1–3]. As bloodmeals are required in every gonotrophic cycle, this has made mosquitoes ideal vectors of blood-borne pathogens such as the malaria parasites.

Inside the mosquito midgut lumen, red blood cell membranes are broken down, and hemoglobin and other proteins of the blood and serum are digested into amino acids, chiefly through the action of bloodmeal-induced trypsin-like and other proteases. Amino acids are then used for protein and fatty acid biosynthesis from reduced carbon atoms [3, 4]. Specifically, carbon skeletons of amino acids are converted to pyruvate, acetyl CoA, or one of TCA cycle intermediates, which in turn are used for energy production through gluconeogenesis or converted to dietary triacylglycerols (TAGs). At the same time, blood-derived TAGs are converted into monoacylglycerols (MAGs) and free fatty acids (FFAs) in the midgut lumen through the action of lipases before they are transported into the midgut cells and converted into diacylglycerides (DAGs), TAGs and phospholipids [1]. TAGs are stored intracellularly within lipid droplets and are transported principally to the fat body from where they provide energy upon demand. TAGs are also synthesized from dietary carbohydrates through glycolysis and gluconeogenesis. The loading and unloading of lipids into and from the fat body is performed by a shuttling mechanism that involves lipophorin (Lp) and other lipid transfer molecules. Lp also caries DAGs from the midgut directly to the ovarian tissues where they convert into TAGs [5]. In addition to their importance as energy resource, lipids are crucial for signal transduction and maintenance of membrane fluidity of the midgut cells [6, 7].

Saturated (SFAs) and unsaturated (UFAs) fatty acids, the building blocks of TAGs and other fats, are carboxylic acids consisting of an aliphatic tail and a terminal carboxyl group. The introduction of the first double bond at the Δ9 position of the aliphatic chain (between carbons 9 and 10) of SFAs, the most critical commitment step in the biosynthesis of mono UFAs (MUFAs), is catalyzed by SCD, an iron-containing, microsomal enzyme conserved across eukaryotes. The reaction requires NADH, cytochrome b5 and molecular oxygen [8]. Major MUFAs such as oleic acid and palmitoleic acid are key precursors of membrane phospholipids, cholesterol esters and triglycerides, and are synthesized from SFAs such as stearic acid and palmitic acid, respectively [8]. The ratio of saturated to unsaturated phospholipids is critical for maintenance of cell membrane fluidity and homeostasis.

SCDs have been investigated for their potential use as chemotherapeutic targets in human metabolic disorders, cancer and infections. For example, the *Mycobacterium tuberculosis* SCD is a known target of the thiourea drug isoxyl [9], and Sterculic oil (SO) is the best recognized natural inhibitor of SCD [10]. The importance of lipid biosynthesis during mosquito blood feeding offers an opportunity for exploitation toward development of novel vector control interventions. Here, we investigate the function of SCD1 in the malaria vector mosquito *Anopheles coluzzii* upon blood feeding. *A. coluzzii* was until recently known as *A. gambiae* M form [11] and is the species used in the majority of experimental studies, most of which still referred to as *A. gambiae*. Silencing *SCD1* through RNA interference and inhibition of SCD1 enzymatic activity by supplementing bloodmeals with SA allowed us to investigate the transcriptional, metabolic and phenotypic impacts of the enzyme on mosquito physiology. The data has led us to conclude that SCD1 is essential for mosquito lipid metabolism, survival and reproduction following a bloodmeal and identify SCD1 as a potential target of novel vector control interventions.

## Methods

### Mosquito colony and maintenance

*A. coluzzii* mosquitoes used in all experiments were of the N’gousso colony established from *A. gambiae* M molecular form and the Forest chromosomal form mosquitoes collected in Cameroon in 2006. Adult mosquitoes were held in netted cages and maintained on 5% fructose solution at 28 °C (±1 °C), 70-80% relative humidity and a 12-hour dark/ light cycle. Larvae were fed on tetraamine fish food.

### Bioinformatics analysis

BLAST searches were conducted using the National Centre for Biotechnology Information (NCBI) and Vector Base sequences and tools. Sequence alignments were performed using locally installed ClustalW and protein prediction was done using the SMART Package (http://smart.embl-heidelberg.de/). Pathway analysis of transcriptomic and metabolomic data was performed using the Kyoto Encyclopedia of Genes and Genomes (KEGG).

### DsRNA preparation and gene silencing

Gene-specific double-stranded RNA (dsRNA) was synthesized and injected as described [12]. Briefly, *SCD1*-specific oligonucleotide primers flanked at their 5’ end by T7 RNA polymerase promoter sequence (*SCD1-RNAi-F*: taatacgactcactataggg-AAATGTGATTGCCTTCGGT; *SCD1-RNAi-R*: taatacgactcactataggg-GCGAGAAGAAGAAGCCAC) were used in PCR reactions on total RNA extracted from 10 sugar-fed adult female mosquitoes using the TRIzol reagent (Invitrogen). DsRNA was synthesized using the T7 MEGAscript kit (Invitrogen) and the PCR product was purified using the RNeasy kit (QIAGEN). DsRNA for the control *LacZ* gene was synthesized as reported [13]. Gene silencing was accomplished by injection of 69 nL (3 ug/uL) of dsRNA into the mesothoracic spiracle of each carbon dioxide-anesthetized 0-2-day old female mosquito. Mosquitoes were allowed to recover for 3 days before any further treatment.

### RNA extraction and qRT PCR

Total RNA was extracted from 10 *dsSCD1* and *dsLacZ* injected mosquitoes, respectively. cDNA synthesis was performed from total RNA following the manufacturer’s instructions for the QuantiTect Reverse Transcription kit (QIAGEN). Gene silencing efficiency was determined using the SYBR *Premix Ex Taq* kit (Takara) according to the manufacturer’s instructions and the ABI Prism 7500 Fast Real-Time thermocycler (Applied Biosystems). The *A. gambiae* ribosomal gene *S7* was used as the endogenous reference, and *SCD1* gene expression was quantified relative to a calibrator *dsLacZ*-injected control samples. qRT primers for *SCD1* were *SCD1-qRT-F:* GGTGTCGAAGGAGATCGTGG and *SCD1-qRT-R:* RTCTGGTTGAGAATGGTGGCG. S7 primers were reported previously [14].

### Transmission electron microscopy

For Transmission Electron Microscopy (TEM), midguts from *dsLacZ* or *dsSCD1* injected *A. gambiae* females were dissected just before blood feeding (0h) and 24h post bloodmeal (PBM) through a standard membrane feeding apparatus (Hemotech). Dissected midguts were fixed for 2h at room temperature (2.5% E.M. Grade Glutaraldehyde, 0.1 M sodium cacodylate buffer containing 2 mM calcium chloride), transferred to 1% osmium tetroxide for 1 h, washed in deionized water, and stained with 2% aqueous uranyl acetate for 30 min. Specimens were dehydrated with ethanol dilutions, infiltrated with Spurr’s resin, and flat-embedded at 60 °C. Transverse sections (70 nm) were prepared using a Leica ultracut UCT ultramicrotome and placed onto 200 thin mesh bar copper grids followed by counterstaining with lead citrate. Midgut sections were viewed with a JEOL-JEM-1230 electron microscope operated at 80 kV and TIFF images (8 bit) were captured using an AMT 4M pixel camera.

### Gene expression profiling

Control (*dsLacZ*-injected) and *SCD1* KD female mosquitoes were fed on human blood though a standard membrane feeding assay 3 days after recovery from dsRNA injection, and 10 fully-engorged female mosquitoes from both groups were sampled at each of the following time points: 0h (just prior to BF), 6h, 12h, 18h and 24h PBM. Total RNA was extracted using the TRIzol reagent (Invitrogen) and purified using the RNeasy kit (Qiagen). RNA quantification was performed using the Nanodrop 1000 spectrophotometer (Thermo Scientific) and RNA quality and integrity was assessed though gel electrophoresis (1% agarose gel) and the Agilent Bioanalyzer 2100 (Agilent Technologies). Labelling and hybridization were performed, following the instructions in the Low Input Quick Amp Labeling kit, for two-color microarray-based expression analysis (Agilent). The Pfalcip_Agamb2009 microarray platform (designed by Dr. Vlachou, personal communication) was used for all of the hybridizations presented. This platform encompasses a total of 15,424 *A. gambiae* annotated transcripts of the AgamP3.8 according to the 2014 VectorBase release along with *P. falciparum* probes. *A. coluzzii* genome assembly suggests 14,712 gene transcripts and shows minor differences from *A. gambiae* Slides were scanned using the Genepix 4000B scanner equipped with Genepix Pro 6.1 software (Axon instruments). Data were normalized using the Genespring 11.0 GX software (Agilent). The Lowess Normalization Method with the threshold of raw signals set to 5 which was enough to eliminate background regulation of *P. falciparum* probes. Significant differences in gene expression between experiment and control groups were evaluated by combining an expression ratio cut-off of 1.0 on a log2 scale and one-way ANOVA statistical tests (with P values of ≤0.025) across different time points. Expression profiles were clustered using the Cluster software, version 3.0, according to the Pearson correlation score. The data were visualized with Java TreeView version 1.1.6r4. All subsequent data analysis was done in Microsoft Excel (Microsoft Corp., Redmond, WA).

### Preparation of SA solution and mosquito treatment

SA was freshly synthesized and kindly provided by Professor Mark Baird in (Bangor University, UK), and 1M stock solution was prepared by dissolving SA in 1M Tris buffer and kept at −20° C. For experiments, SA stock solution was diluted further into Tris buffer and human blood to a final concentration of 1mM. Four-day old female mosquitoes were fed on human blood supplemented with a 1mM SA (final concentration in blood) in 1mM Tris buffer or with 1mM Tris buffer alone. Mosquitoes were maintained on 5% sucrose solution throughout the experiments. For survival analysis, dead mosquitoes from each group were counted every 6h post bloodmeal (PMB). Before performing experiments with 1mM SA concentration, different concentration of SA including 0.4, 0.6, 0.8, 1, 1.4, 1.6 and 1.8 mM were tested to determine the IC50.

### Metabolomic Profiling

*SCD1* KD and *dsLacZ*-injected (control), and SA-treated and 1mM Tris-treated (control) mosquitoes were sampled at 0h (just prior to blood feeding), 18h and 36h PBM. Sample preparation for metabolomic profiling and data analysis were performed as previously described [15]. In brief, 3 mosquitoes, from each time point, were extracted in a 1 ml ice-cold solution of methanol: water (8:2 v/v) and centrifuged (14000 rpm, 4° C, 15 min) to separate the cell debris from the supernatant. The supernatant was then transferred to a silanized 1.5 ml glass vial (Agilent Technologies UK Ltd) and dried in a SpeedVac concentrator (Eppendorf). Methoxymation, in combination with trimethylsilylation (methoxy-TMS), was accomplished for derivatization. Sample analysis was performed on an Agilent 7890 GC coupled to a 5975c mass spectrometer, using the Fiehnlib settings, running a pooled biological quality control (pbQC) sample every six samples [16]. Myristic-d27 acid was used as the endogenous reference. Deconvolution and integration of the extracted metabolites was performed using an in-house work-flow as described [17]. All experiments were carried out in 5 independent biological replicates with 3 technical replicates per each biological replicate. To produce heat maps, metabolite data was normalized to the internal standard (Myristic-d27 acid) and pbQC levels, then the averages of each metabolite from five replicates for control or treated group at different time points were calculated. Finally, each metabolite was calibrated to the highest value for that metabolite across all time points and groups. Significant differences in metabolite signals between experimental and control groups were evaluated by combining metabolite ratio cut-off 0.1 on a log2 scale. MATLAB was used to plot the data.

### Antibiotic treatment

To reduce the natural microbiota load in the midgut, mosquitoes were treated with antibiotic solutions as described previously [13]. In brief, mosquitoes were collected just after emergence and kept on a cocktail of 25 μg/ml gentamycin, 10 μg/ml penicillin and 10 units/ml streptomycin, diluted in 5% sucrose. The treatment continued for 5 days, with the antibiotic solution refreshed every 24 hours. Total RNA was extracted from 10 *dsSCD1* and *dsLacZ*-treated mosquitoes per replicate, followed by cDNA synthesis from total RNA was performed using the QuantiTect Reverse Transcription kit (Qiagen). 16S rDNA primers used were as reported previously [13].

## Results and Discussion

### SCD1 is essential for adult female survival upon blood feeding

Bioinformatics searches of the *A. gambiae* and *A. coluzzii* genomes identified *AGAP001713* as a putative stearoyl-CoA Δ9-desaturase, designated as *SCD1*. The gene encodes a 355 amino acid-long protein with a single transmembrane domain and a putative FA desaturase domain (68-309 aa) showing 45.1%, 92.2% and 71.9% identity to equivalent domains of *SCD* homologues in human, *Drosophila melanogaster* and *Plasmodium falciparum*, respectively (**Fig. S1**). The desaturase domain contains three Histidine (His) boxes designated as region Ia, Ib and II, which together contain 8 conserved His residues that are essential for SCD1 activity.

Newly emerged, age-matched (0-2-days old) female mosquitoes were injected with control *LacZ* and *SCD1* dsRNAs, respectively, kept separately in netted cups and maintained on 5% w/v glucose solution. *SCD1* silencing efficiency 3 days post recovery was 78% as assessed by qRT-PCR (**Fig. 1A**). At day 3 post recovery, the mosquitoes were fed on human blood through a membrane feeder. Fully engorged mosquitoes were separated, counted and maintained on 5% w/v glucose solution throughout the rest of the experiment. Dead mosquitoes from each group were counted every 6 hours post bloodmeal (PBM). The results revealed a dramatic increase of mortality in *SCD1* KD blood-fed mosquitoes starting at about 24h PBM and reaching 100% by 48h PBM (**Fig. 1B**). The median survival time was 42h. Only 17% mortality was observed in control mosquitoes at 48h PBM, which was significantly higher than that of *SCD1* KD mosquitoes (p<0.0001).

**Fig. 1.**
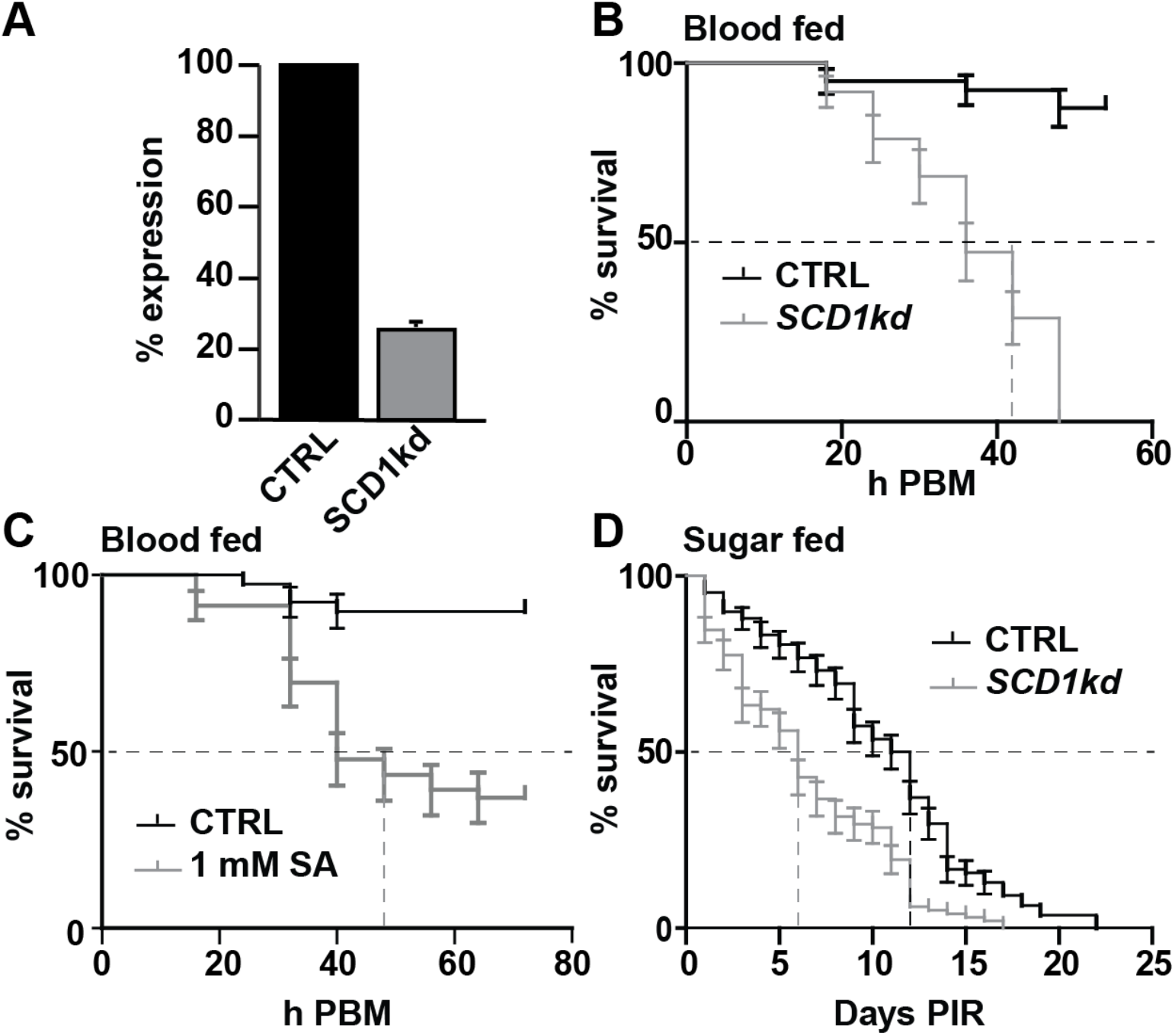
Blood and sugar meal induced mortality in *SCD1* KD and SA treated female *A. coluzzii*. (**A**) Efficiency of RNAi-mediated silencing of *SCD1*. Silencing was measured with qRT-PCR 3 days post recovery following dsRNA injection of freshly hatched adult females. Control (CTRL) mosquitoes were injected with dsRNA against the *LacZ* gene. (**B-D**) Survival analysis of adult female mosquitoes that were: (B) injected with *SCD1* or control (CTRL) *LacZ* dsRNAs following feeding on human blood (post blood feeding; PBF) at day 3 post injection, (C) treated with 1mM SA or control Tris buffer, added to human blood provided through a membrane feeder, or (D) injected with *SCD1* KD or CTRL *LacZ* dsRNA and maintained on 5% sucrose for 25 days post injection recovery that started at day 3 post injection (PIR). Survival was recorded every 6 h. Data were analyzed using Kaplan-Meier survival analysis and log-rank statistics (Mantel-Cox test), showing highly statistical differences between treatments and controls in all three sets (P<0.0001). The mean ± SEM of data obtained from 3 independent biological experiments and the median survival time (dashed lines) are shown.

Survival assays were repeated, this time following supplementation of the human blood with 1 mM SA in 1 mM Tris buffer instead of *SCD1* silencing. Human blood supplemented with 1 mM Tris buffer alone was used as a control. Similar to the *SCD1* KD assays, mosquitoes fed on blood supplemented with SA also exhibited increased mortality rates compared to controls, albeit not as rapid, which reached 52% at 42h PBM (p<0.0001; **Fig. 1C**). Interestingly, the mortality rate was reduced after 42 h, presumably due to quick SA break down.

We examined whether the observed survival phenotype was specific to blood-fed mosquitoes by monitoring the survival of *SCD1* KD and *dsLacZ* control mosquitoes that were not blood-fed but maintained on 5% sucrose solution throughout the experiment. Whilst mortality rates of *SCD1* KD sugar-fed mosquitoes were evidently slower than the *SCD1* KD blood-fed mosquitoes, these rates were significantly higher than their sugar-fed controls (p<0.0001; **Fig.1D**). The median survival time for the *SCD1* KD group was 6 days and for the control group was 11.5 days.

### *SCD1* silencing affects midgut epithelial integrity and egg development

Light microscopy 24h PBM revealed a blood-filled thoracic cavity in *SCD1* KD mosquitoes compared to controls, indicating blood perfusion of the hemocoel compartment (**Fig. 2A, top panel**). In addition, *SCD1* KD midguts were extremely fragile and appeared swollen, with bloodmeal not contained within a well-defined blood bolus (**Fig. 2A, middle panel**). These phenotypes suggested that *SCD1* KD may affect the midgut physiology and cell membrane fluidity, directly or indirectly compromising the ability of mosquitoes to cope with the mechanical and metabolic stress of blood feeding.

**Fig. 2.**
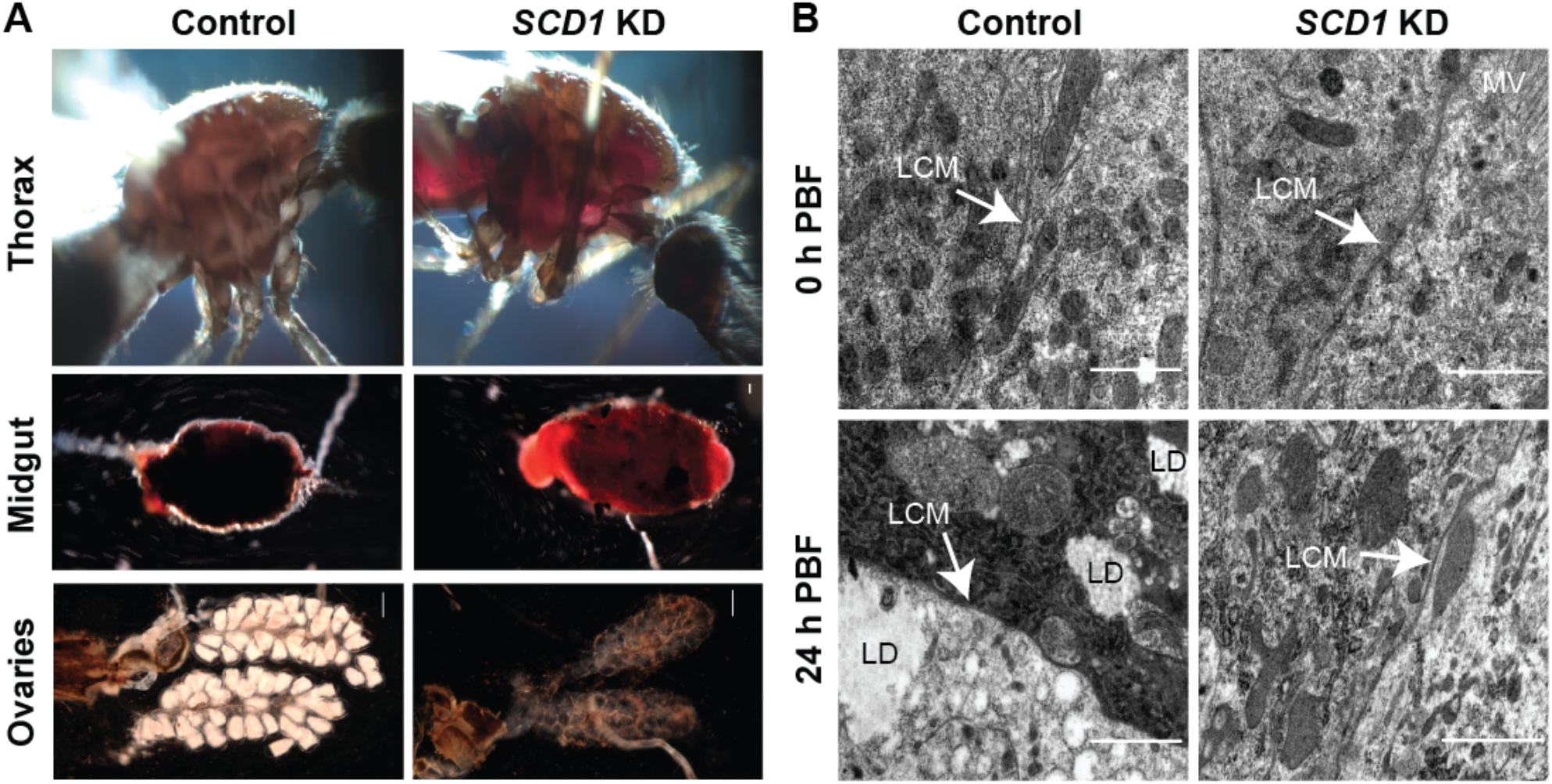
Phenotypic characterization of *SCD1* KD mosquitoes. (**A**) Representative pictures of thoraces, midguts and ovaries of blood-fed control (left) and *SCD1* KD (right) female mosquitoes at 24h PBM. (**B**) Representative TEM pictures of midgut epithelial cells dissected from control (left) and SCD1 KD (right) female mosquitoes at 0h before blood meal and 24h PBM, respectively. LCM denotes lateral borders between adjacent midgut cells. Thickness of LCM is the mean value of 10 different points on the same LCM. LD denotes lipid droplets and MV denotes microvilli. Scale bar is set at 1 μm.

We investigated the subcellular basis of the *SCD1* KD phenotype using transmission electron microscopy (TEM) on midgut tissues at 0h (just before bloodmeal) and 24h PBM. Imaging revealed that midgut cell membranes were thicker in *SCD1* KD (mean width 60 nm) compared to control (31nm) mosquitoes prior to bloodmeal (**Fig. 2B, top panel**), suggesting loss of cell membrane rigidity in agreement with previously reported induction of negative cell membrane curvature [18, 19]. At 24h PBM, cell membranes were substantially thicker in *SCD1* KD (55 nm) than control (14 nm) midguts. Noteworthily, cell membrane thickness in *SCD1* KD midguts was only marginally reduced after bloodmeal (60 nm compared to 55 nm respectively). Reduction of cell membrane thickness is a known adjustment to the expansion of the epithelial cell surface in order to provide structural support to the engorged midgut epithelium [20]. We conclude that *SCD1* loss-of-function alters the ability of midgut epithelial cells to undergo adjustments needed to offset the increased pressure within the lumen, causing rapturing of the epithelial cell layer upon blood feeding and blood perfusion of the thoracic cavity.

TEM analysis also revealed a great deficiency in lipid droplets in *SCD1 KD* compared to control midgut cells at 24h PBM (**Fig. 2B, bottom panel**). Therefore, blockage of SCD1-catalyzed MUFA synthesis appears to inhibit TAG and phospholipid biosynthesis from bloodmeal-derived carbon in the midgut cells as reported previously [4]. These data prompted us to examine the effect of *SCD1* KD on egg development. Ovaries are a major target tissue of Lp-carried lipids synthesized in the midgut from bloodmeal-derived carbon. Indeed, *SCD1* KD mosquitoes showed no signs of egg maturation following a bloodmeal (**Fig. 2A, bottom panel**), similar to the phenotype previously observed upon silencing Apolipophorin II/I (ApoII/I), the main component of the insect Lp [21]. We conclude that compromised lipid synthesis in the midgut cells due to inhibition of SCD1 is responsible for reduced lipid shuttling in the hemolymph leading to underdeveloped ovaries.

### *SCD1* silencing greatly impacts on metabolic homeostasis

We investigated the molecular basis of the *SCD1* KD phenotype in female mosquitoes in response to blood feeding using DNA microarray expression profiling of *SCD1* KD versus *dsŁacZ*-injected control mosquitoes. Five time points from three independent biological replicates were examined: 0h (just prior to blood feeding), 6h, 12h, 18h and 24h PBM. In total, 1432 genes registered expression values in at least one of the time points examined. Genes with at least 2-fold expression changes (p<0.05 in ANOVA t-test) in *SCD1* KD versus control mosquitoes, in any of the time points, were analyzed further. To derive a general picture of expression changes, genes were functionally classified into 9 major functional categories as done previously [21]. Overall, upregulation dominated downregulation in most of the time points; however, there was a marked difference between the different categories (**Fig. 3** and **Table S1**). *SCD1* KD considerably disturbed metabolic homeostasis that is essential for mosquito survival and reproduction, as well as other gene functional categories, all of which are presented in this section as they provide an integrated picture of the disruption of mosquito physiology. The impact of *SCD1* KD on the immune response is analyzed later.

**Fig. 3.**
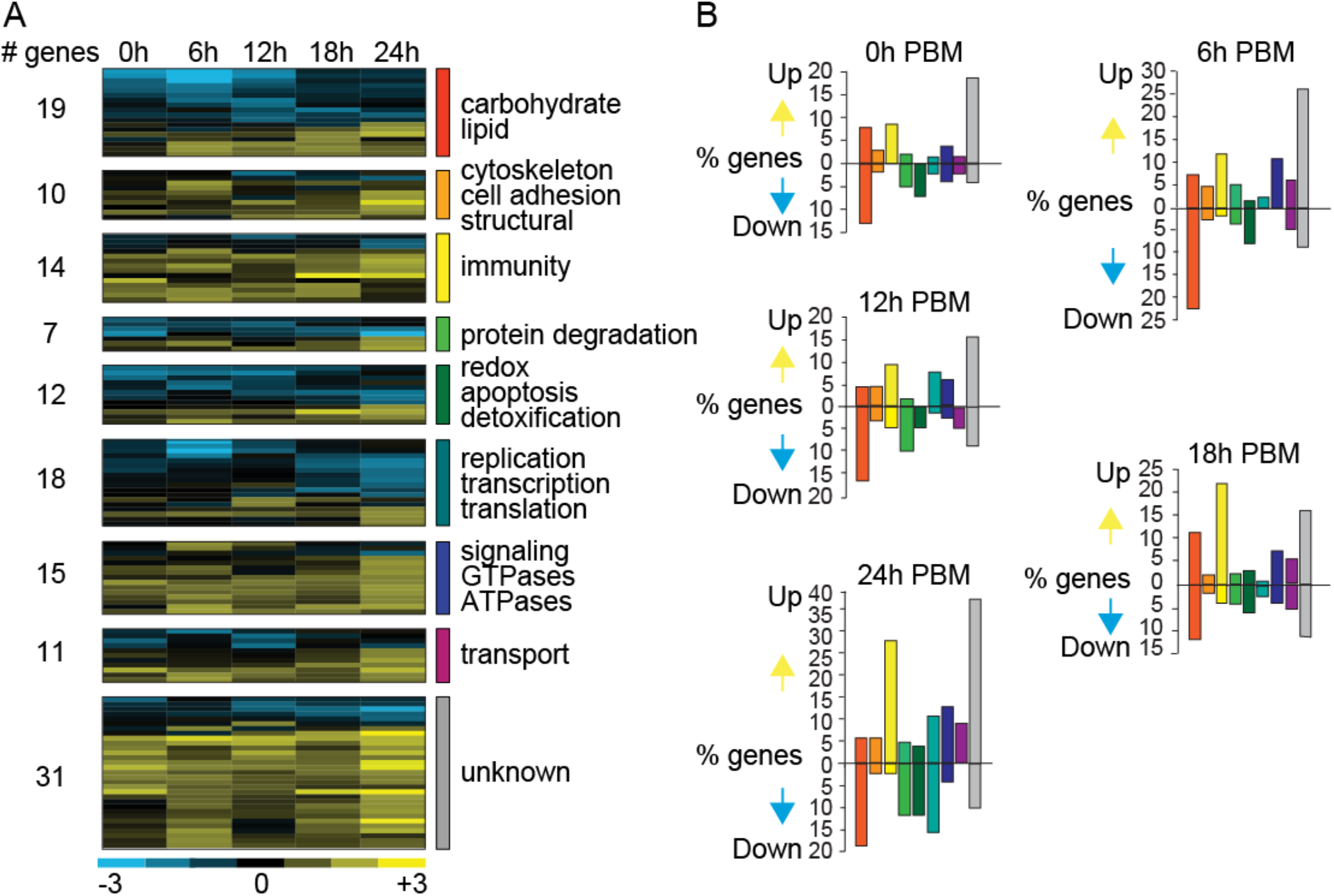
Gene transcription profiles upon blood feeding of *SCD1* KD mosquitoes. (**A**) Heat map showing the transcription profiles of a total of 137 genes exhibiting at least 2-fold differential expression in *SCD1* KD vs. *dsLacZ*-injected control across all time points. The time points examined are 0h (just prior to blood meal), 6h, 12h, 18h and 24h post bloodmeal (PBM). Blue color indicates downregulation and yellow color indicates upregulation, as shown in the color bar below the heat map. The data shown is the mean from 3 independent experiments. The number of genes grouped into each of 8 functional categories or are of unknown function is shown. (**B**) The graphs show the percentage of genes that are upregulated or downregulated in each time point and belonging to each of the 8 functional categories or having unknown function (grey colored bars). The color code of bars representing each of the functional categories is the same as the color code shown in panel A.

#### Carbohydrate and lipid metabolism

A total of 79 genes involved in “*carbohydrate and lipid metabolism*” were differentially regulated in at least one time point; of these, 73% were downregulated and 19 were differentially regulated across the time course. Approximately 39% and 29% of all the genes differentially regulated at 6h and 24h PBM belonged to this category.

Differential expression of lipid metabolism genes was noticed (21/28), with several genes involved in lipid biosynthesis (6/10), lipolysis (5/6) and lipid transportation (4/9) being downregulated. Genes encoding acetyl-CoA carboxylases (ACC) that catalyze the irreversible conversion of acetyl-CoA to malonyl-CoA were also downregulated. As silencing *SCD1* increases SFA accumulation by inhibiting SFA to MUFA conversion, it is hypothesized that SFA buildup over time hinders ACC activity through a well-known feedback mechanism resulting in a drop of malonyl-CoA and increased transport of acetyl-CoA into the mitochondria for β-oxidation [22]. 3-hydroxy-3-methylglutaryl-CoA lyase (HMG-CoA lyase) that releases acetyl CoA by breaking down leucine and fats [23]was downregulated across all time points. We speculate that HMG-CoA lyase downregulation is due to the increased flux of acetyl-CoA caused by over-activation of β-oxidation. UDP glucuronosyltransferase (UGT) that catalyzes glucuronidation removing toxic compounds such as fatty acid derivatives from body was also downregulated. Insect UGTs play important roles in cuticle formation, pigmentation and olfaction, besides their major role in inactivation and excretion of endogenous and exogenous compounds [24, 25]. This may reflect a feedback mechanism to prevent the removal of fatty acids.

Importantly, Lp was markedly downregulated (>3 fold, at 6h and 12h PBM) in *SCD1* KD mosquitoes, which may highlight a feedback regulatory mechanism caused by the depletion of lipid droplets in the midgut cells due to the blockade of SFA to MUFA conversion. Silencing ApoII/I is previously shown to inhibit egg development, which is consistent with the *SCD1* KD phenotype reported here [26, 27]. Surprisingly, phospholipid scramblase (PLS1), a plasma membrane-anchored enzyme that shuttles phospholipids between the inner and outer leaflets of the cell membrane to maintain membrane integrity, was upregulated (>2 folds, at 6h PBM). This response may represent an effort to sustain the SFA/ MUFA ratio in *SCD1* KD cells.

Genes encoding TCA cycle enzymes including isocitrate dehydrogenase (IDH) that catalyzes the reversible conversion of isocitrate to alpha-ketoglutarate, and succinate-CoA synthetase (SUCS-GDP forming) and succinate-CoA synthetase (SUCS-ADP forming), which catalyze the conversion of succinate and GTP or ATP to succinyl-CoA and ADP or GDP, respectively, were also downregulated at 6h PBM. This suggests higher succinyl-CoA involvement in the TCA cycle [28] in *SCD1* KD mosquitoes. Limitation of free CoA in *SCD1* KD mosquitoes may also be an important factor for the slowdown of the forward reaction of TCA cycle and accumulation of TCA cycle intermediates.

A large number of carbohydrate metabolism genes (41) including many involved in glycolysis and gluconeogenesis (11) as well as pyruvate/acetyl-CoA metabolism (8) were differentially regulated in *SCD1* KD mosquitoes. Interestingly, α-glucosidase (AGL) and α-amylase (AAM), which convert glycogen/starch and their intermediates to glucose were downregulated (>2 fold at 0h, 6h and 18h PBM), which could imply lower levels of glucose and other downstream metabolites of glucose. However, downregulation of two key gluconeogenesis enzymes, namely pyruvate carboxylase (PC) and phosphoenol pyruvate carboxykinase (PEPCK), was also observed. At the same time, two key enzymes of the Leloir pathway, lactate dehydrogenase (LDH) and galactokinase (GALK), were upregulated (>3 fold at 6h and >2 fold at 0h PBM, respectively). Since the Leloir pathway converts lactate/galactose to glucose/glucose derivatives in order to maintain homeostasis during glucose scarcity, it is possible that activation of this pathway together with PC and PEPCK upregulation is part of an effort to compensate for glucose deficiency.

#### Protein metabolism

As many as 46 genes involved in the “*protein degradation*” category was differentially regulated in *SCD1* KD compared to control mosquitoes, of which 67% were downregulated, mostly at 24h PBM. Seven (7) of these genes were differentially regulated across the experimental time course. Mosquitoes require a variety of proteolytic enzymes to digest the bloodmeal. Among downregulated genes were several exopeptidases, including alanine aminopeptidase, serine protease S28, astacin, carboxypeptidase A1 and various trypsins. Nevertheless, some proteolytic enzymes were upregulated including peptidase M19 (6h PBM), peptidase S33 (12h PBM), trypsin 4 (18h PBM) and peptidase S60 (24h PBM). It is reported that enzymes expressed early after a bloodmeal such as trypsin 4 may indirectly activate the transcription of the major endoproteolytic enzyme trypsin 1 [29]. Taken together, these data suggest that a complex network of proteolytic processes involved in bloodmeal metabolism are differentially affected in *SCD1* KD mosquitoes contributing to the dramatic phenotype observed.

Arginosuccinate synthase (ASS) and arginosuccinate lyase (ASL), two key enzymatic components of the arginine biosynthetic pathway, whereby citrulline is converted to arginine releasing fumarate, were downregulated in *SCD1* KD mosquitoes (18h and 24h PBM). It is plausible that accumulation of fumarate leads to inhibition of ASS and ASL expression [30]. The upregulation of ornithine decarboxylase (ODC), which converts ornithine to putrescine may suggest further metabolism of putrescine initially to spermidine and then through the glutathione detoxification pathways.

#### Cytoskeleton, cell adhesion and structural components

Several genes (27) belonging to the “*cytoskeleton, cell adhesion and structural components*” category showed differential expression throughout the time course; the majority of them were upregulated. Importantly, upregulation of an actin-like gene was detected at 6h PBM. It was previously reported that increased levels of actin mRNA at 3-4h PBM correlates with epithelial remodeling to accommodate distention of the midgut upon blood ingestion [20]. As *SCD1* KD causes rigidity of the epithelial cell membranes, this is suggestive of a stress response to overcome the mechanical pressure caused by the rigid cell membranes. E-Cadherin, a Calcium-dependent cell adhesion protein involved in cell-cell interactions was previously shown to be overexpressed in *SCD1* silenced breast cancer cells [31].Consistent with this finding, an upregulation of a gene encoding a cadherin-like protein was observed 24h PBM. This protein is expressed in the midgut of *A. gambiae* larvae, where it plays a key role in detoxification of the nonchemical larvicide-Bti [32, 33]. Thus, its overexpression could serve in detoxifying excess SFAs.

A gene encoding a chitin-binding domain protein of the peritrophic matrix (PM) was consistently downregulated throughout the time course but 18h PBM, with a peak >5-fold decrease at 24h PBM. *A. gambiae* PM is detectable as early as 12h PBM and fully formed by 48h PBM [34]. We hypothesize that PM synthesis is compromised in *SCD1* KD mosquitoes owing to the downregulation of this gene. Indeed, the juvenile hormone (JH) binding protein (JHBP) was also upregulated at 24h PBM. JH plays important roles in larval development, reproduction and stress management and is known to suppress PM synthesis and function in adult *Calliphora erythocephala* [35]. As JH levels drop with increasing JHBP levels [36, 37], JHBP upregulation in *SCD1* KD mosquitoes may play part in suppression of ovary development and PM synthesis.

#### Signaling/ATPases/GTPases

JHBP belongs to the “*signaling/ATPase/GTPase*” gene category. Of this category, 52 genes in total were differentially regulated throughout the time course, mostly at 6h and 24h PBM representing a quarter and a third of all upregulated genes at these time points, respectively; the majority of them upregulated and 15 of them differentially regulated across the time course.

Insulin-like peptides (ILPs) that can have a multitude of functions including lipid and glycogen processing, immunity, reproduction and longevity [38, 39] were significantly downregulated across 5 time points. In *Ae. aegypti*, ILP3 is known to stimulate ovaries to synthesize ecdysteroid hormones including 2-hydroxyecdysone that induces production of vitellogenin in the fat body [40, 41]. A possible reduction of ecdysteroids could also lead to lower levels of JH [36, 42]. These data could suggest interruption of insulin mediated signaling in *SCD1* KD mosquitoes and perhaps defective ecdysteroid hormone synthesis.

The insulin/mTOR (mammalian target of rapamycin) signaling plays a central role in regulating cell growth and proliferation by sensing and integrating a variety of inputs arising from amino acids, cellular stresses, energy status and growth factors [40]. The phosphatidylinositol 5-phosphate 4-kinase (PIP4K), one of the essential mediators of many membrane signaling events and a modulator of mTOR Complex 1 (mTORC1) activity in insects was upregulated. Overexpression of PIP4K in *Drosophila* upregulates the activity of mTORC1 and increases the levels of mTOR downstream [43, 44]. We hypothesize that the increased levels of amino acids and substrates of TCA cycle in response to blood feeding in *SCD1* KD mosquitoes, induces the expression of PIP4K to enhance the mTORC1 function [40].

The gene encoding the alpha subunit of the Na/K-transporting ATPase, which catalyzes the cation transfer across the membrane by converting ATP to ADP, was highly upregulated at 6h, 12h and 24h PBM further suggesting disruption of the epithelial cell membrane integrity and homeostasis.

Genes encoding odorant binding proteins OBP13 and OBP26 were markedly upregulated across all 5 time points in *SCD1* KD mosquitoes. OBPs are shown to have strong affinity for SFAs like palmitate, stearate, octanoic acid and hexanoic acid [45]. Therefore, the accumulation of SFAs in *SCD1* KD mosquitoes may trigger overexpression of these OBPs.

#### Replication, transcription and translation

Significant differential regulation was observed at 24h PBM for 64% of genes (29/ 48) in this category, with 35% of them (18/48) exhibiting differential expression across all 5 time points. Nuclear export factor RNA Binding Motif protein 15 (RBM15) that is required for efficient mRNA export from the nucleus [46] was consistently downregulated in *SCD1* KD mosquitoes across the time course. Cyclin A, an important regulator of cell cycle progression, was also downregulated at 24h PBM. The link between SCD1 and cell cycle regulators is largely unknown. However, recent data indicate that SCD1 blockage reduces AKT activity by dephosphorylation, with subsequent activation of glycogen synthase-kinase (GSK3-β), which in turn degrades cyclin D1 thereby inhibiting cell proliferation. AKT is a key component of the survival signaling pathway, and GSK3β is a downstream element of AKT signaling that is regulated by insulin [47]. Together these data suggest that *SCD1* KD disrupts cell cycle progression in *A. coluzzii*.

#### Redox, apoptosis and detoxification

A total of 37 genes in this category were differentially regulated in the *SCD1* KD mosquitoes. The majority of them (83%) were downregulated, mostly at 24h PBM, and a third of them (12/37) were differentially regulated across all time points.

The detoxification gene *GstD7*, which plays an important role in protecting against cellular damage by controlling oxidative stress [48, 49], was consistently upregulated in *SCD1* KD mosquitoes across all time points. *GstD7* is highly expressed in *Drosophila suzukii* upon exposure to the pesticide malathion [50]. Cytochrome c oxidase polypeptide 7A (COX7A), the terminal component of the mitochondrial respiratory chain that catalyzes the reduction of oxygen to water was downregulated across all time points. COX7A is shown to be downregulated during acute inflammation [51], supporting our hypothesis of a potent inflammatory response in *SCD1* KD mosquitoes.

Genes encoding cytochrome P450s (CY314A1, CY302A1 CYP12F1, CYP6P1 & CYP9K1) were significantly downregulated at different time points. Previous studies demonstrate that cytochrome P450s play significant role in lipid homeostasis by assisting lipid degradation [52]. Hence, it is possible that scarcity of TAGs leads to the downregulation of cytochrome P450s as a feedback response. Glutathione-S-transferase D7 (GSTD7), a major detoxifying enzyme in mosquitoes, was upregulated, which may be due to the SFA:MUFA ratio distortion and represent a homeostatic response to hinder lipid peroxidation.

#### Transport/vesicle mediated transport

As many as 26 genes involved in transport/ vesicle mediated transport system were differentially regulated in *SCD1* KD compared to control mosquitoes, of which 15 were upregulated, mostly at 24h PBM. Trehalose transporter 1 (TRET1) that transports trehalose disaccharide from the fat body to hemolymph [53] was downregulated across the time course. Silencing trehalose transporter is shown to reduce trehalose content in hemolymph and shorten the lifespan of *A. gambiae* under stress conditions like desiccation and high temperature [54, 55].

### Metabolomic profile of *SCD1* KD mosquitoes

We used GC-MS-based metabolomic analysis of deproteinized extracts to examine the metabolic changes induced in female mosquitoes prior to (0h PBM) and upon feeding on human blood (18h and 36h PBM) following *SCD1* silencing or SA (in 1 mM Tris) treatment. Mosquitoes injected with *dsLacZ* or treated with 1 mM Tris were used as controls, respectively. Differences in metabolite signals between experimental and control groups were evaluated.

Desaturase indices, the FFA product to substrate, serves as a biomarker for the desaturase activity. The results showed that *SCD1* KD causes a significant reduction in desaturase indices 16:1/16:0 (palmitoleate to palmitate) at 18h and 36h PBM (**Fig. 4A**) and 18:1/18:0 (oleate to stearate) at all time points (**Fig. 4B**). A statistically significant reduction in both 16:1/16:0 and 18:1/18:0 indices was also observed upon SA treatment albeit only at 36h PBM. These data confirm that silencing *SCD1* or inhibition of the desaturase activity blocks the conversion of SFAs to UFAs. They are also in agreement with other studies correlating decreased SCD1 activity with increased SFA content, caused by administration of the cyclopropenoic fatty acids SA and malvalic acid [56].

**Fig. 4.**
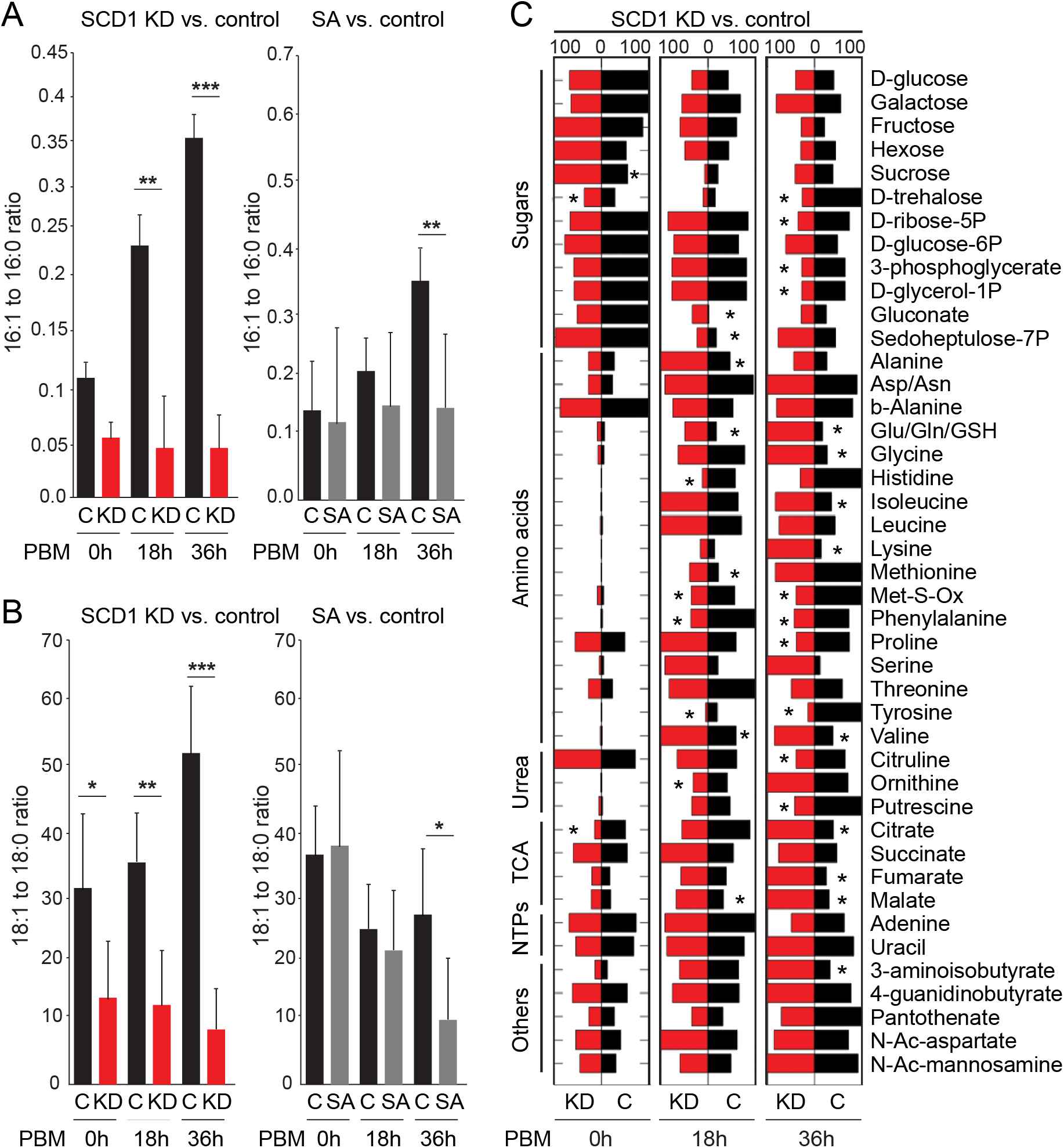
Metabolic responses in blood fed SCD1 KD and SA treated mosquitoes. Desaturase indices (**A**) 16:1/16:0 (Palmitoleate to Palmitate) and (**B**) 18:1/18:0 (Oleate to Stearate) in *SCD1* KD (left) and SA-treated (right) adult female mosquitoes prior to (0h) and 18h and 26h post bloodmeal (PBM). *DsLacZ* injected and 1mM Tris treated mosquitoes were used as controls for *SCD1* KD and SA-treatment, respectively. Reported values are mean ± SE of 5 independent biological replicates, where n=3 mosquitoes per group per replicate. *p<0.05, *p<0.001, ***p< 0.0001. (**C**) Metabolite profiles in *SCD1* KD and SA-treated mosquitoes and their respective *dsLacZ* and 1mM Tris treated controls. Only metabolites which showed statistically significant (*p<0.01) changes in the percentage of metabolite signals in the experimental group compared to the control group in at least one of the conditions are shown in the heat map. Red bars represent the mean percentage of metabolite signals in *SCD1* KD or SA treated mosquitoes from 5 independent biological repeats. Black bars show mean percentage of metabolite signals in respective controls.

The metabolomic analysis of 43 aqueous metabolites showed significant differences (P<0.01 or less) of several metabolite concentrations between the *SCD1* KD compared to control mosquitoes at one or more of the examined time points (**Fig. 4C**). In *SCD1* KD mosquitoes, the levels of substrates of the TCA cycle including citrate (36h PBM), malate (18h and 36h PBM) and fumarate (36h PBM) increased in response to blood feeding, while the intermediates of glycolysis, glycerol-3P (also 36h PBM), and the pentose phosphate pathway including ribose 5-P (36h PBM) decreased in response to blood feeding. The levels of D-glucose and D-glucose-6P remained unchanged.

Female mosquitoes largely convert protein-rich liquid meals into amino acids after blood ingestion. Consistent with this, only traces of most amino acids were observed at 0h PBM which significantly increased after bloodmeal in *SCD1* KD mosquitoes at 36h PBM compared to that of the control. During oogenesis, phenylalanine is preferentially channeled towards late protein synthesis in order to meet the need for egg cuticle hardening in *A. gambiae* [15]. Interestingly, decreased levels of phenylalanine and – even more so – of its derivative tyrosine were observed in *SCD1* KD mosquitoes at 18h and 36h PBM. It has been previously shown that interference with *A. gambiae* phenylalanine hydroxylase resulting in reduced tyrosine formation leads to fewer eggs and retarded vitellogenesis [15]. It can be therefore inferred that *SCD1* KD leads to a perturbed conversion of phenylalanine to tyrosine and a subsequent suppression of egg development. Indeed, Decreased levels of histidine may be linked to the role of histidine as a major intracellular buffering agent in animal cells [57]. Buffering agents maintain cellular pH and little changes in pH can cause either acidosis or alkalosis in the body with deleterious effects on metabolism. Hence, decreased levels of histidine may indicate imbalance in the pH system of *SCD1* KD mosquitoes. Increased concentration of essential amino acids, such as isoleucine, valine, glutamic acid/glutamine, alanine, lysine and methionine may be the result of a feedback mechanism to improve amino acid uptake.

The levels of ornithine, a central component of the urea cycle, were higher in *SCD1* KD at 18h and 36h PBM compared to control whereas the levels of putrescine were increased at 36h PBM.

### Integrative metabolic pathway analysis

To better understand metabolic perturbations caused by *SCD1* KD, 17 genes and 21 metabolites from the transcriptomic and metabolomic analyses, respectively, were mapped onto cellular metabolic processes (**Fig. 5**). Pathway analysis provided insights into *SCD1* KD-induced cellular toxicity dynamics in mosquitoes and revealed that silencing *SCD1* significantly disrupts the TCA cycle and its interwind pathways with increasing accumulation of substrates and feedback regulation of expression of metabolic enzymes.

**Fig. 5.**
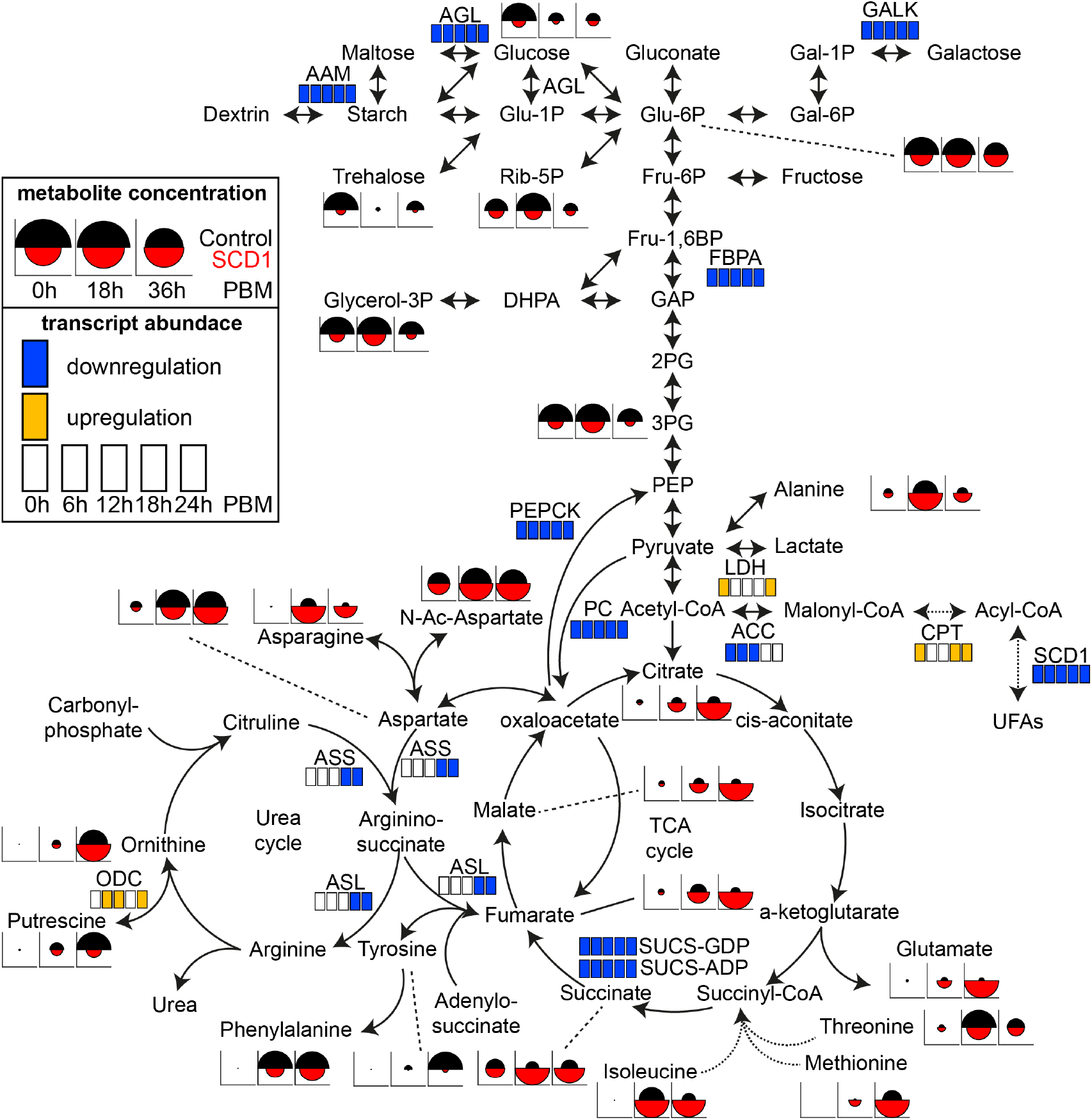
Integrative analysis of metabolic pathways. Metabolite and transcript abundance differences between ds*SCD1* (KD) and ds*LaZ* (control) injected mosquitoes are mapped on reconstructed KEGG pathways encompassing glycolysis, gluconeogenesis, TCA cycle and Urea cycle. Transcript differences in any of five time points are indicated as upregulated (yellow) or downregulated (blue) when 2-fold or more expression changes are documented in *SCD1* KD versus control mosquitoes (p<0.05 in ANOVA t-test). The metabolite data were normalized for each metabolite basis and plotted as circle fragments of which the radius is directly proportional to the highest value for the metabolite in any of the three time points and two conditions. AGL, α-glucosidase; AAM, α-amylase; PC, pyruvate carboxylase; PEPCK, phosphoenol pyruvate carboxykinase; LDH, lactate dehydrogenase; GALK, galactokinase; LDH, lactate dehydrogenase; FBPA, fructose-bisphosphate aldolase; SUCL-GDP, succinate-CoA ligase; SUCS-ADP, succinate-CoA synthetase; ACC, acetyl-CoA carboxylases; ASS, arginosuccinate synthase; ASL, arginosuccinate lyase; ODC, ornithine decarboxylase; CPT, carnitine-palmitoyl transferase; Glu-6P, glucose 6-phospate; Glu-1P, glucose 1-phospate; Gal-6P, galactose 6-phospate; Gal-1P, galactose 1-phospate; Rib-5P, ribose 5-phospate; PEP; phosphoenol pyruvate; Fru-6P, Fructose 6-phospate; Fru-1,6BP, fructose 1,6-biphospate; GAP, glyceraldehyde-3-phosphate; 2PG, 2-phosphoglycerate; 3PG, 2-phosphoglycerate; PEP, phosphoenol pyruvate.

Silencing *SCD1* results an increased accumulation of SFA over time by modulating SFA/MUFA ratios that hinder acetyl-CoA carboxylases (ACC) activity through a well-known feedback mechanism resulting in a drop of malonyl-CoA that down-regulates carnitine-palmitoyl transferase (CPT) and in turn increases transport of acetyl-CoA into mitochondria for β-oxidation [58]. This upregulated β-oxidation in turn increases the reaction rate of many of the steps in the TCA cycle, thereby increasing flux throughout the pathway and resulting in higher accretion of TCA cycle intermediates over time, i.e. citrate, succinate, fumarate and malate. The significantly higher levels of amino acids that can fuel the TCA cycle at various steps, such as methionine, isoleucine and valine, which is thought to be part of a feedback mechanism to increase amino acid uptake could indicate a major cause of accretion of these metabolites in *SCD1* KD mosquitoes. Therefore, accumulation of TCA cycle intermediates is the likely cause of the decreased expression of rate limiting enzymes such as SUCS-GDP forming and SUCS-ADP forming enzymes, as well as those that feed into the TCA cycle including arginosuccinate synthase (ASS) and arginosuccinate lyase (ASL) observed in *SCD1* KD mosquitoes in order to achieve homeostasis. The much higher levels of asparagine and glutamate may also be part of the feedback mechanism to suppress the TCA cycle through depletion of intermediate metabolites.

The low intracellular levels of the polyamine putrescine, produced through the urea cycle, in *SCD1* KD compared to control mosquitoes could be a direct result of the low ASS and ASL levels. However, it may also be due to the parallel upregulation of ornithine decarboxylase (ODC), which converts ornithine to putrescine, and suggest increased metabolism of putrescine initially to spermidine, which is also low in *SCD1* KD mosquitoes (not shown), and then though the glutathione detoxification pathways. Hence the levels of ornithine appear to be unaffected between SCD1 KD and control mosquitoes.

Lower levels of sugar and downstream substrates of the glucose metabolism, such as ribose-5P (Rib-5P, which controls the rate-limiting purine synthesis), trehalose, 3-phosphoglycerate (3PG) and – to a lesser extent – glucose-6P (Glu-6P), in *SCD1* KD mosquitoes compared to controls may indicate suppression of gluconeogenesis as a feedback mechanism, also suggested by the strong PEPCK downregulation. Interestingly, *SCD1* KD mosquitoes appear to activate compensatory pathways, such as the Leloir gluconeogenic pathway, by upregulating the galactokinase (GALK) and lactate dehydrogenase (LDH) to replenish the glucose flux. Downregulation of fructose-bisphosphate aldolase (FBPA) that converts fructose 1, 6-BP to glyceraldehyde-3-phosphate (GAP), both precursors of dihydroxyacetone phosphate (DHAP), cause low levels of glycerol-3P which would lead to reduced glycerol-based phospholipids, the main component of biological membranes.

In conclusion, the metabolic pathway analysis suggests that inhibition of *SCD1* in female mosquitoes reprograms the interwinding pathways for cellular metabolism, leading to accumulation of various metabolites causing cytotoxicity and death.

### *SCD1* KD triggers a robust immune response

Immunity-related genes became increasingly and proportionally more numerous in the *SCD1* KD upregulated gene set as time progressed (**Fig. 3** and **Table S2**). A total of 58 genes were significantly differentially regulated in at least one of the time points, 83% of them upregulated and 14 genes differentially regulated across the time course. Immune induction was particularly prominent at 18h and 24h PBM.

One fifth of differentially regulated genes (19%) encode clip-domain serine proteases and serine protease homologs, including CLIPA5, CLIPA8, CLIPB11-12, CLIPB14-20 and CLIPB20; all but CLIPB20 upregulated. Many of these proteins were previously shown to be involved in microbial lysis and melanization in the mosquito hemolymph [59–62], suggesting activation of a systemic immune response in *SCD1* KD mosquitoes. The melanization catalyzing enzyme PPO3 was also upregulated at 24h PBM.

TEP3, a member of the thioester-containing protein family, was amongst upregulated genes. TEP3 is paralogous to the main complement pathway effector, TEP1, but does not have a thioester motif and therefore is presumed to have a regulatory rather than effector function. Like TEP1, TEP3 binds to the putative receptor/adaptor complex LRIM1/APL1C and is involved in responses against bacteria and malaria parasites [63, 64]. Another component of the complement pathway upregulated in *SCD1* KD mosquitoes was CTLMA2, a member of the C-type lectin protein family. CTLMA2 is transcriptionally induced in response to infection and is thought to promote pathogen lysis and/or phagocytosis and negatively regulate melanization [65].

Fibrinogen related proteins, FREPs, play key roles in mosquito humoral and systemic responses and are thought to have complementary and synergistic functions, some seemingly acting as pattern recognition receptors (PRRs) [66]. Strong upregulation in *SCD1* KD mosquitoes was observed for *FREP19, FREP21, FREP24, FREP32, FREP59* and *ficolin A*. Another putative PRR, the scavenger receptor SCRBQ3, was also upregulated in *SCD1* KD mosquitoes. SCRBQ3 is involved in melanization and induced in response to injury and associated opportunistic infections as well as upon injection of Sephadex beads [67, 68].

The lipid-binding MD2-like proteins (MLs), ML4 and ML7, were strongly upregulated at 18h PBM in *SCD1* KD. MLs are thought to bind lipids, and the founding member of the family, MD2, synergizes with TLR4 to trigger expression of pro-inflammatory cytokines in response to bacterial lipopolysaccharide (LPS). We hypothesize that mosquitoes upregulate ML4 and ML7 in response to abnormal and excessive saturated fatty acids, activating systemic reactions. Indeed, SFAs are shown to be recognized by the TLR4-MD2 complex, triggering trigger inflammatory pathways, similar to LPS [69].

There are two possible hypotheses for the observed upregulation of immune-related genes, most of which are involved in systemic immune reactions of the hemolymph, in *SCD1* KD mosquitoes. The first is that over-activation of the immune system is caused by the gut microbiota (or their immune inducers) that gain increased access to immune receptors on the midgut epithelium or the hemolymph due to the loss of the peritrophic membrane and epithelium integrity. The second hypothesis is that the increased concentrations of SFAs and/or purinergic molecules such as ATP and ADP trigger an auto-inflammatory signaling. In invertebrates, humoral substances, often called danger associated molecular patterns (DAMPs), released by damaged cells or tissues, activate signaling pathways leading to inflammation.

We investigated the microbiota-induced response hypothesis by examining the expression of the antimicrobial peptide gene *CEC1*, which was upregulated across all time points in the microarray experiments, in *SCD1* KD and control (ds*LacZ*-injected) mosquitoes that were treated with antibiotics provided though the sugar meal since adult emergence to rid their gut from resident microbiota. As a control, mosquitoes were fed only on sugar. After 5 days, mosquitoes were either sampled (0 h) or fed on human blood though a membrane feeder and sampled 24 hours later. The load of bacteria in the mosquito gut was quantified by qPCR of the 16S rRNA at 0h PBM. Thee independent biological replicates were performed.

The results showed that *CEC1* was significantly upregulated both at 0h PBM and, more so, at 24h PBM in *SCD1* KD compared to control mosquitoes (**Fig. 6**). Surprisingly, the microbiota load prior to bloodmeal was significantly lower in SCD1 KD mosquitoes compared to controls. These results indicate that induction of *CEC1* expression in SCD1 KD mosquitoes is independent of the microbiota load, which is decreased in *SCD1* KD mosquitoes either due to immune activation or due to the physiological and metabolic conditions in the gut. Indeed, a much stronger upregulation in *SCD1* KD mosquitoes was detected in mosquitoes treated with antibiotics. As expected, the bacterial load was substantially lower in antibiotic-treated compared to non-treated mosquitoes at 0h PBM, and even lower in *SCD1* KD antibiotic-treated mosquitoes. These data reject the hypothesis that over-activation of the immune system in *SCD1* KD mosquitoes is caused by the gut microbiota and suggest that these mosquitoes may suffer from an auto-inflammatory condition caused by high concentrations of SFAs, purinergic molecules and/or TCA cycle intermediates. They additionally suggest that the microbiota may be playing an inhibitory role to immune signaling in *SCD1* KD mosquitoes, possibly triggered by the distorted SFA:MUFA ratio and mediated though microbiota tolerance pathways.

**Fig. 6.**
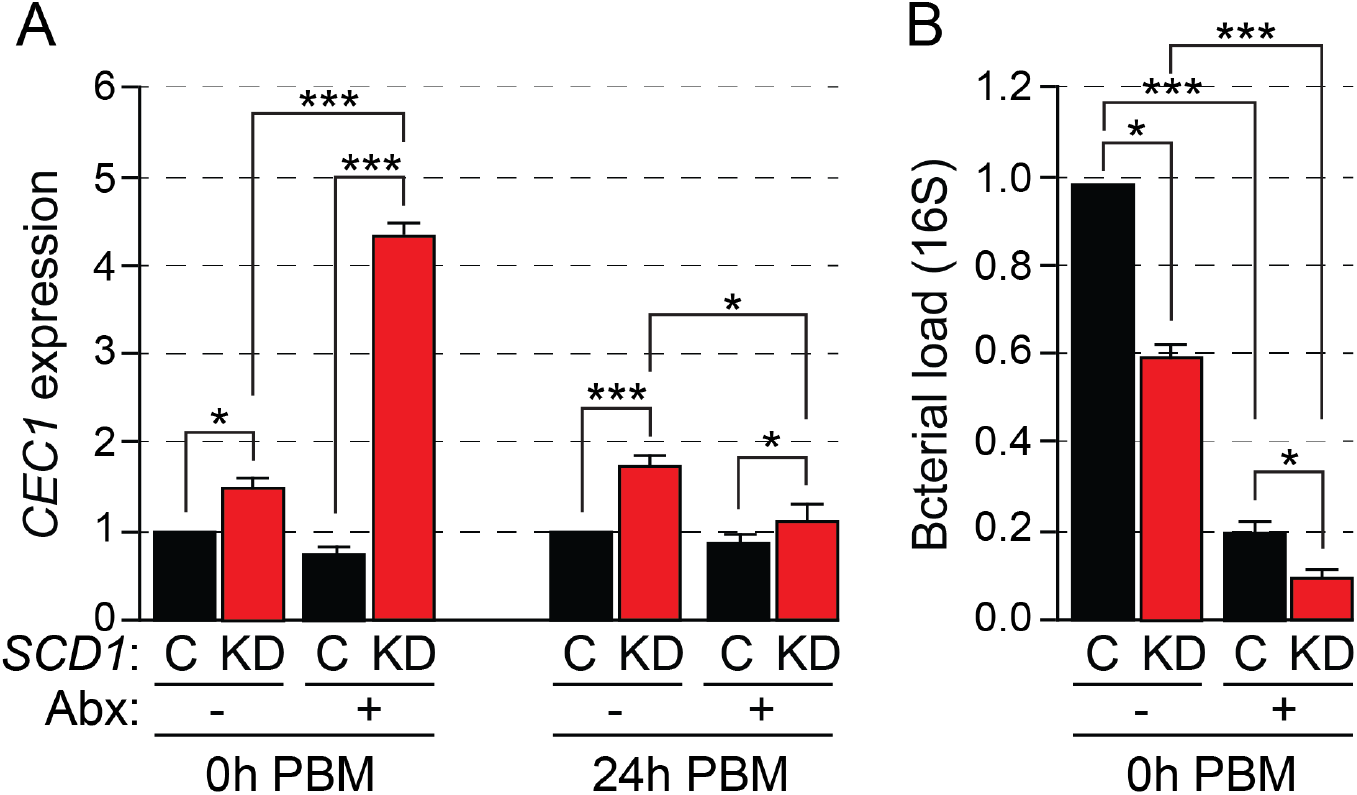
Impact of gut microbiota on immune activation in *SCD1* KD mosquitoes. (**A**) Relative expression of the AMP *CEC1* expression in dsLacZ-injected (C) and SCD1 KD (KD) adult mosquitoes fed on sugar (-) or sugar plus antibiotic (Abx; +) solution since emergence. After five, mosquitoes were sampled either without further treatment (0h PBM) or after they have been provided a blood meal (24h PBM). (**B**) Bacterial load in the midguts of mosquitoes treated as above, measured by qRT-PCR of 16S rRNA. Reported values are the mean ± SE of three independent biological experiments of 10 mosquitoes per group. ***p< 0.0001, *p<0.01.

## Conclusion

Stearoyl-CoA desaturase is a rate limiting key enzyme regulating the ratio of SFAs to UFAs that is essential in the maintenance of cell membrane fluidity, proliferation, lipid accumulation and signaling. Its loss of function is known to alter lipid homeostasis due to the overloading with SFAs, inducing a cellular toxic response known as lipotoxicity, which eventually modulates the global metabolic profile and leads to various physiological abnormalities in animal cells [4]. Here we demonstrate that loss-of-function of SCD1 in adult female *A. coluzzii* mosquitoes upon blood feeding, the main vector of human malaria in West sub-Saharan Africa, leads to a highly skewed SFA:UFA ratio, loss of cell membrane integrity, transcriptional and metabolic reprogramming and various other detrimental physiological effects, including an acute auto-inflammatory condition and compromised reproduction. We therefore conclude that SCD1 is a good target of interventions to control vector abundance and block malaria transmission, including small molecule inhibitors (insecticides and drugs) and transgenesis methods aiming at mosquito population suppression or replacement.

## Acknowledgements

We thank Professor Mark S. Baird for kindly providing us with Sterculic Acid and Katarzyna Sala for her assistance with mosquito rearing. Z.F. was supported by a Commonwealth Scholarship. The work was supported by a Wellcome Trust Project Grant 093587/Z/10/Z to G.K.C. and D.V. and a Wellcome Trust Investigator Award 107983/Z/15/Z to G.K.C.

---

**Fig. S1.**
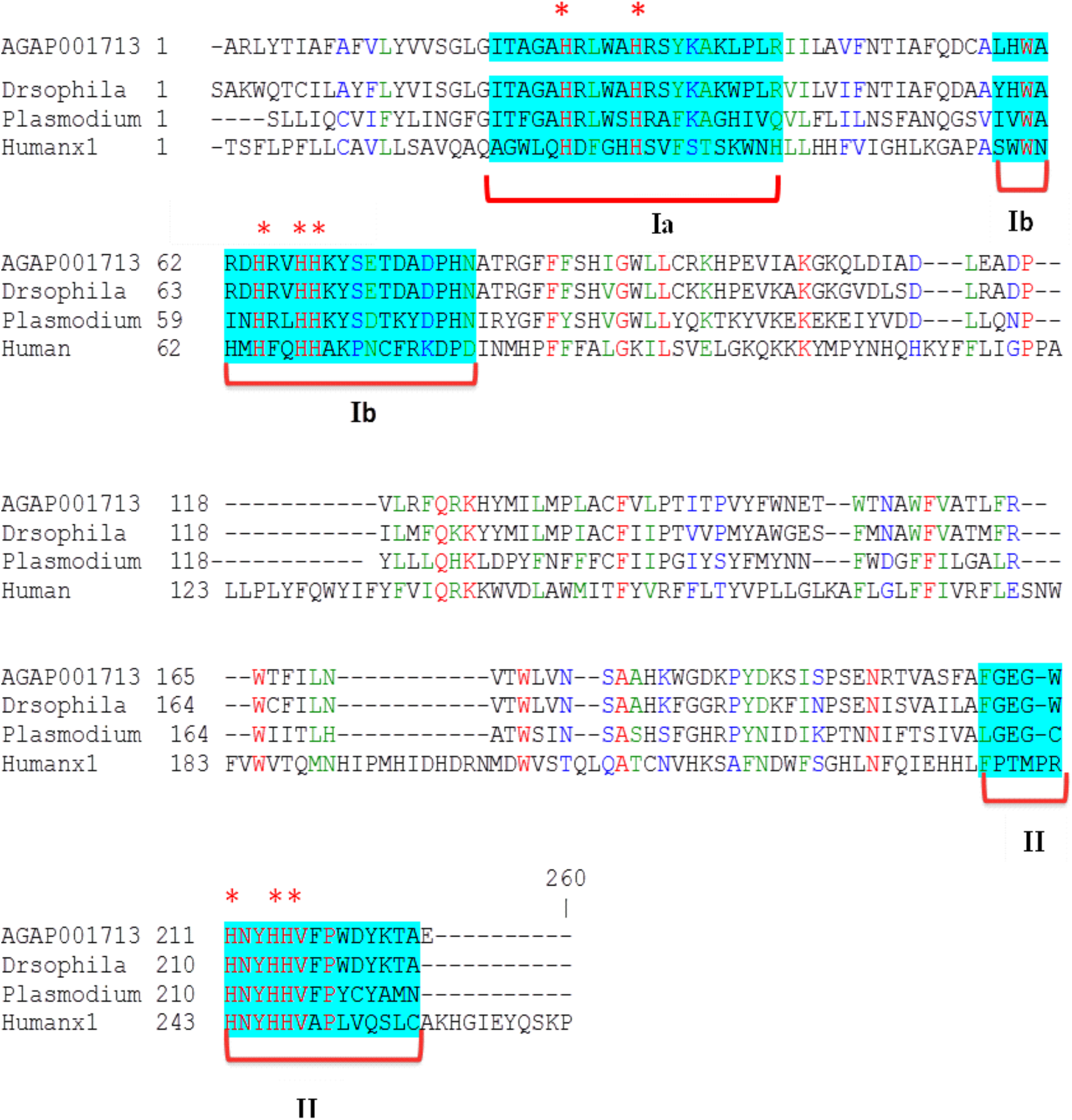
Sequence analysis of the *A. coluzzii* Stearoyl-CoA desaturase1. Alignment of the central domain of SCD1 (amino acid residues 88 to 303) and the equivalent domain of SCD1 orthologues in human, *Drosophila melanogaster* and *Plasmodium falciparum*, performed using CLUTALW. The three His boxes designated as region Ia, Ib and II, respectively, and the eight conserved His residues are shown.

**Table S1.**
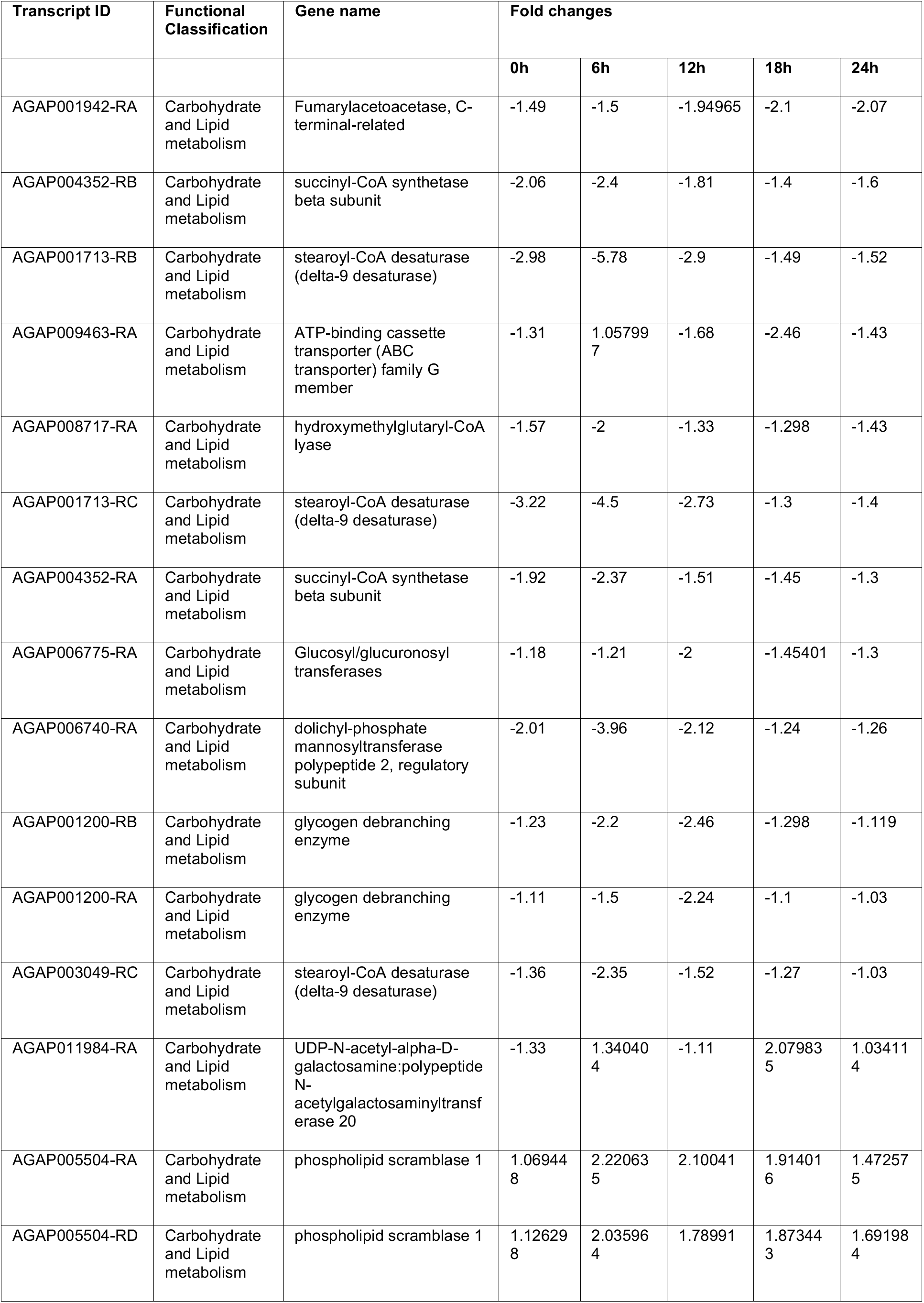
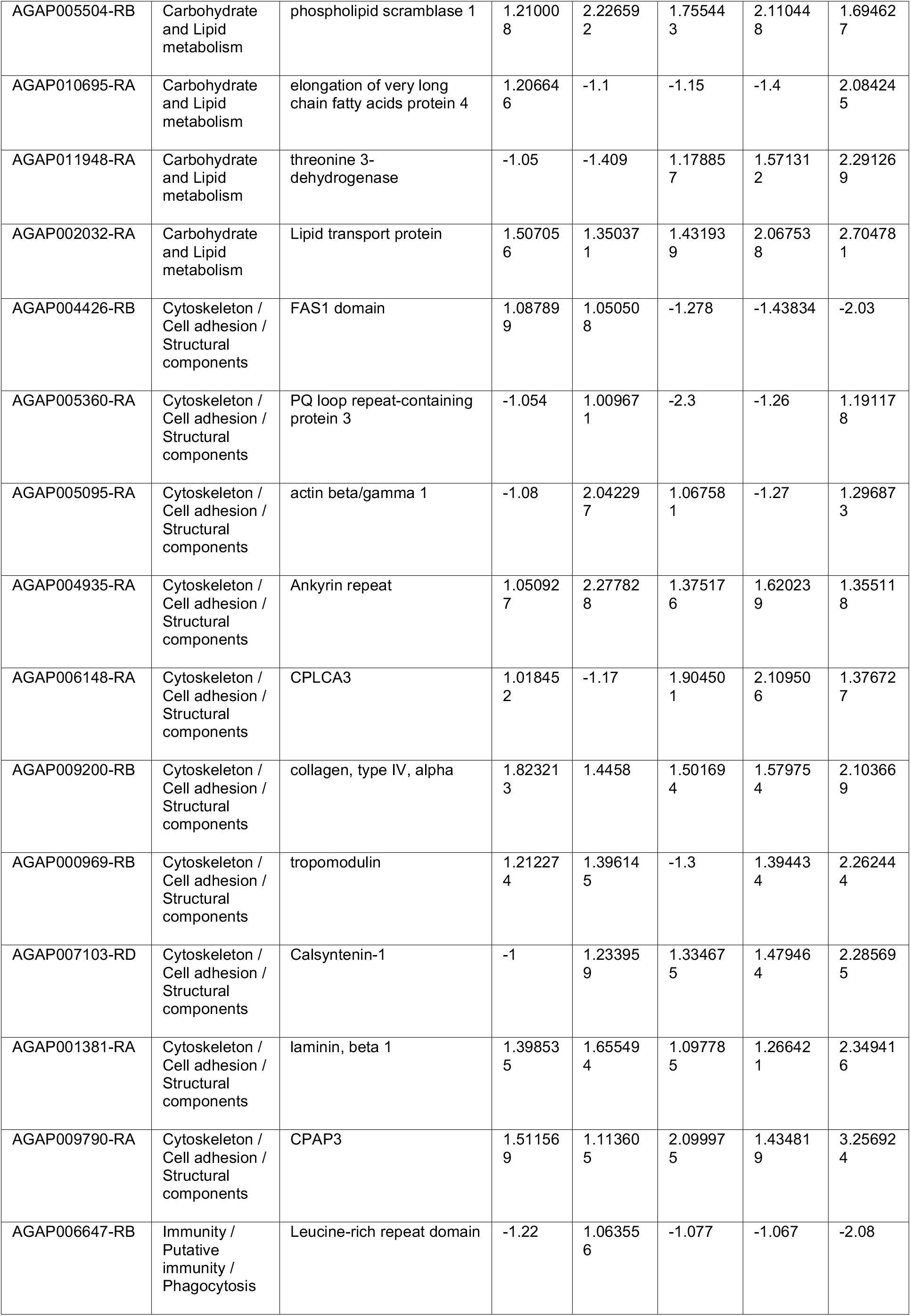
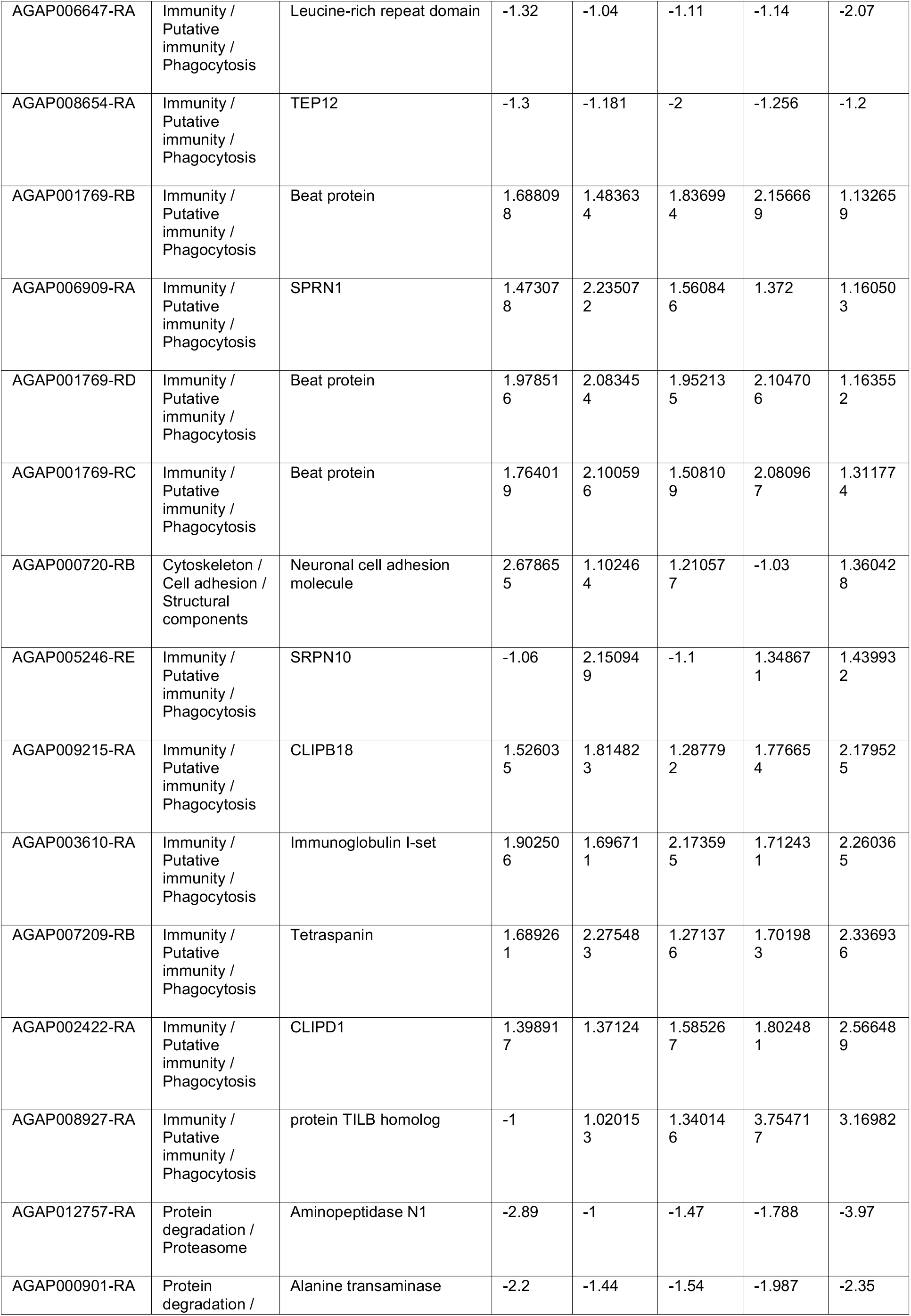
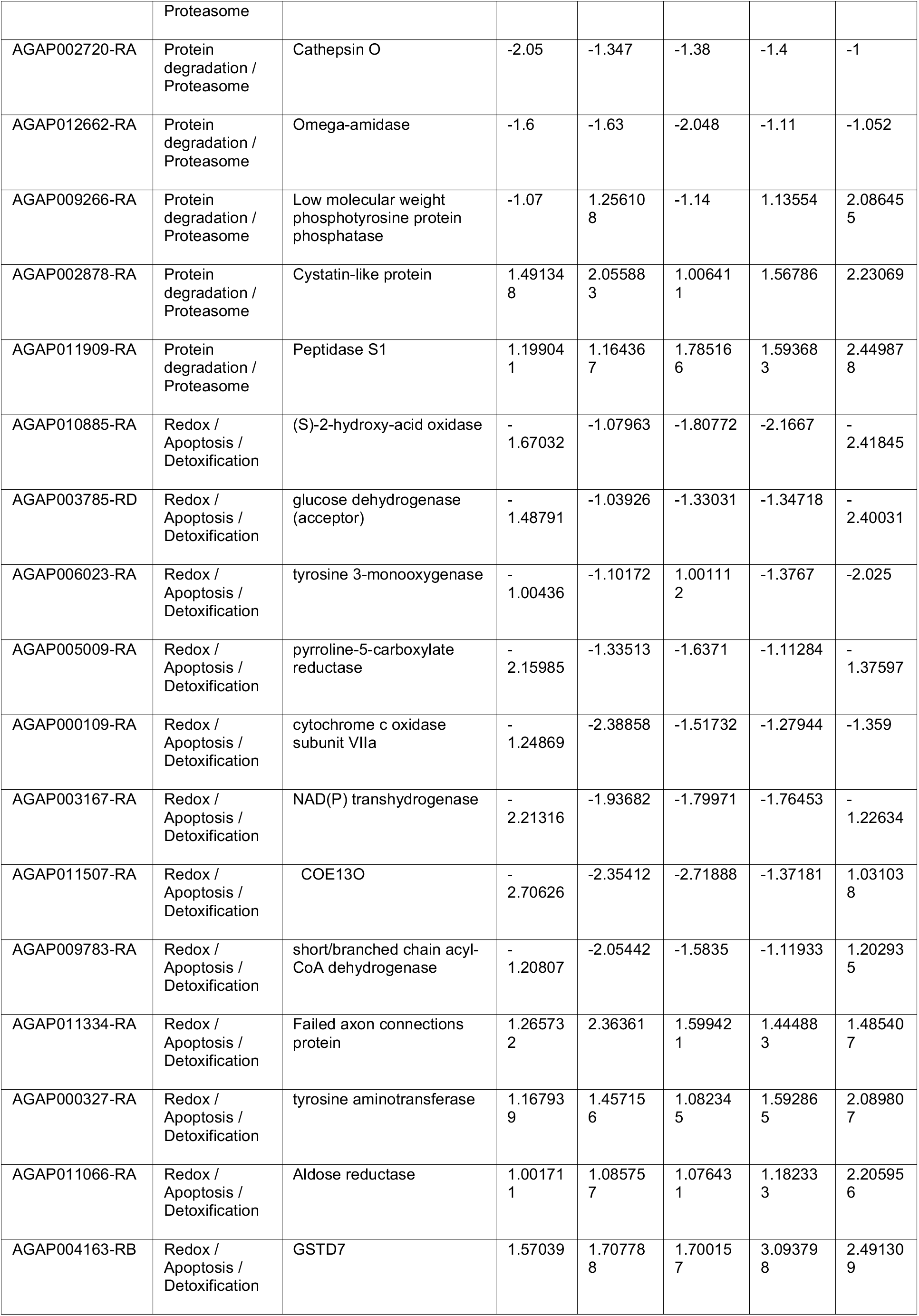
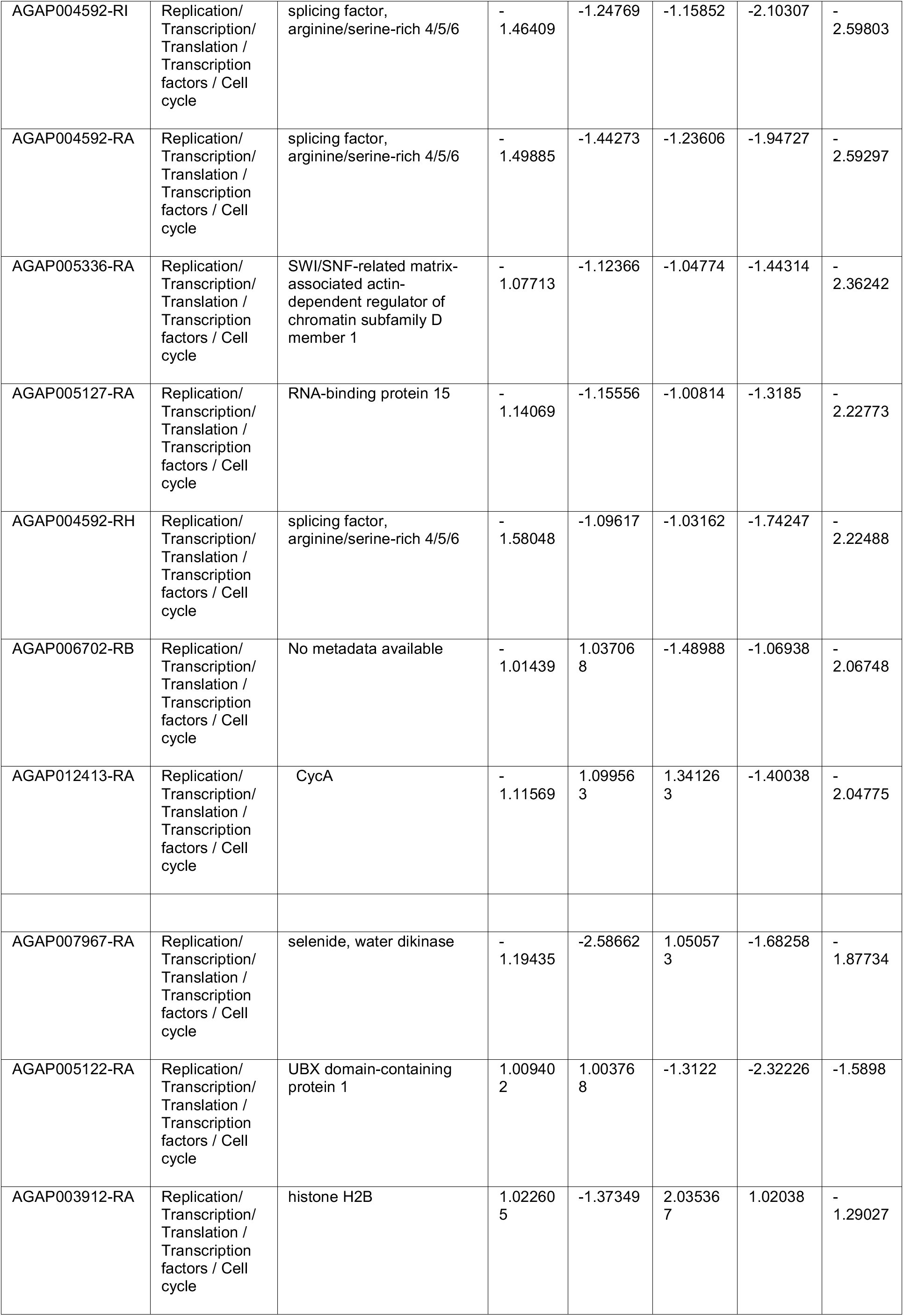
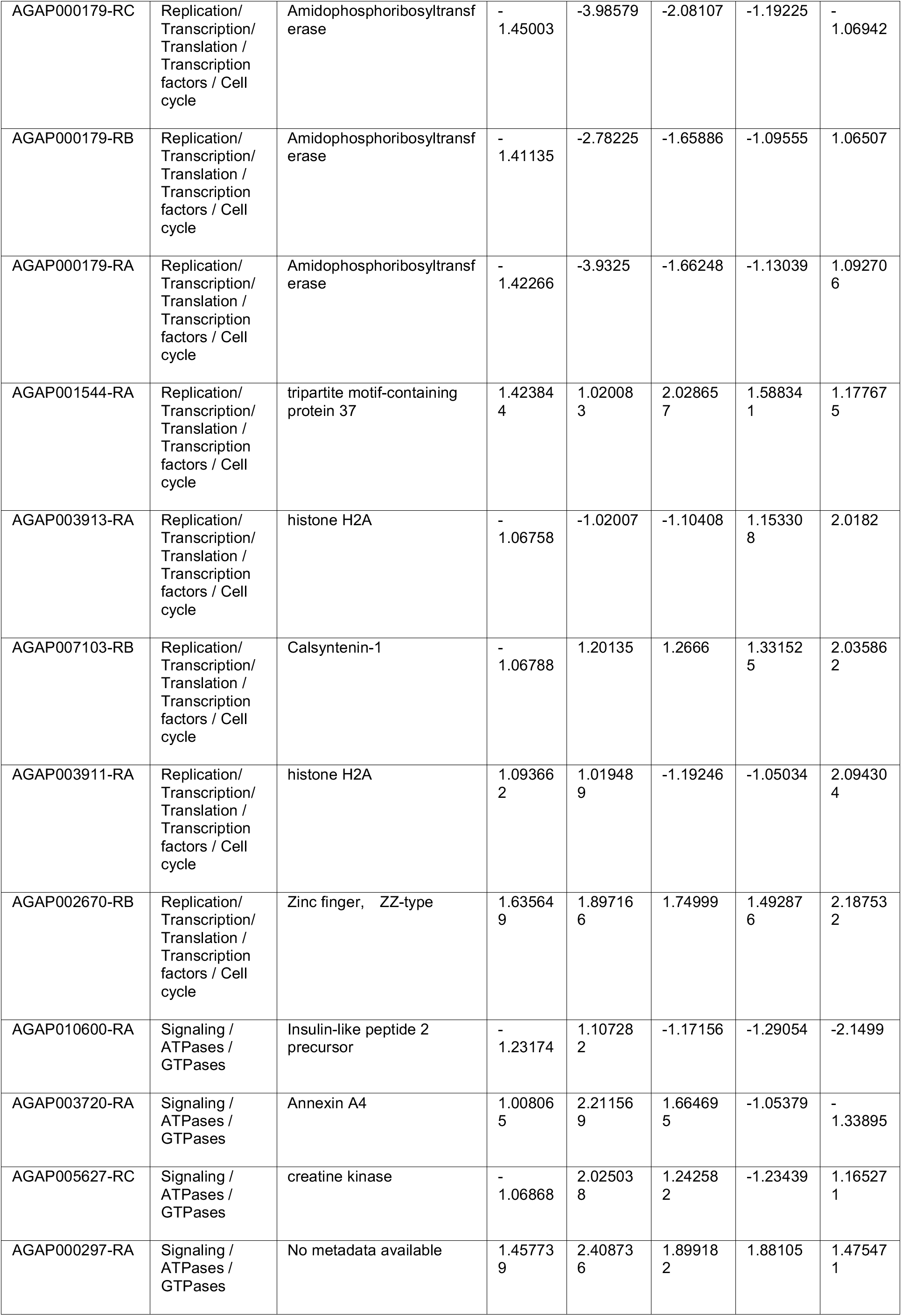
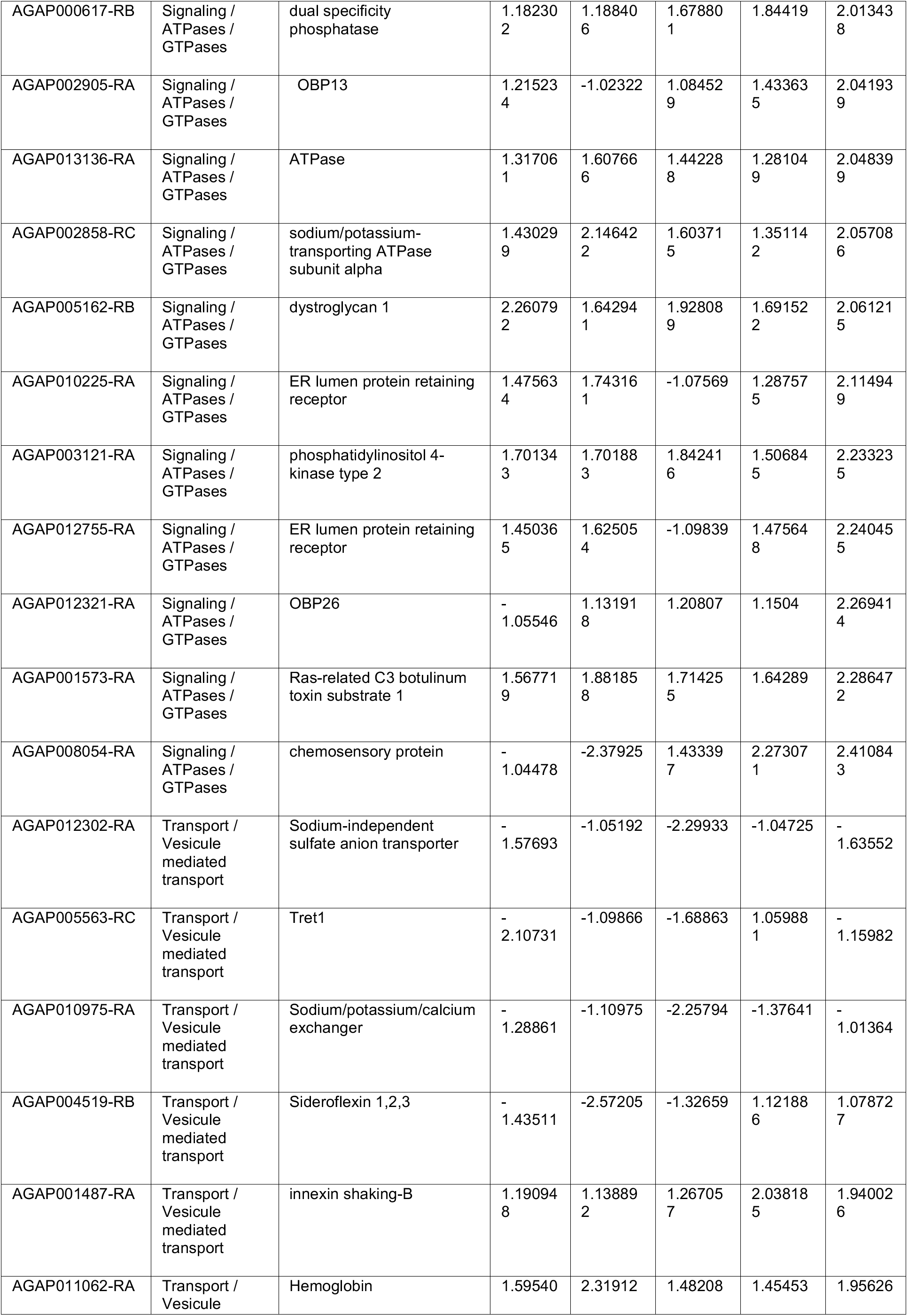
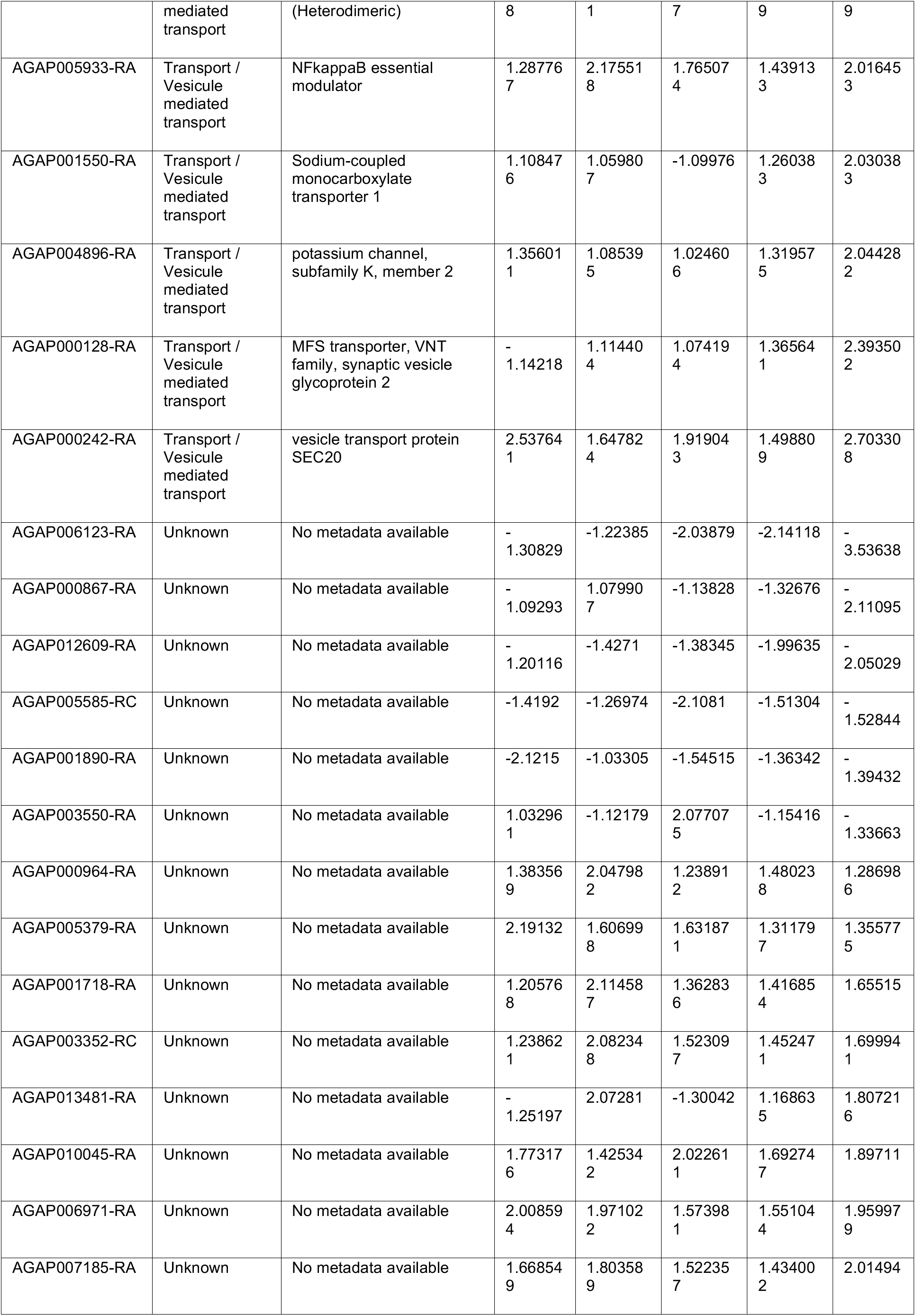
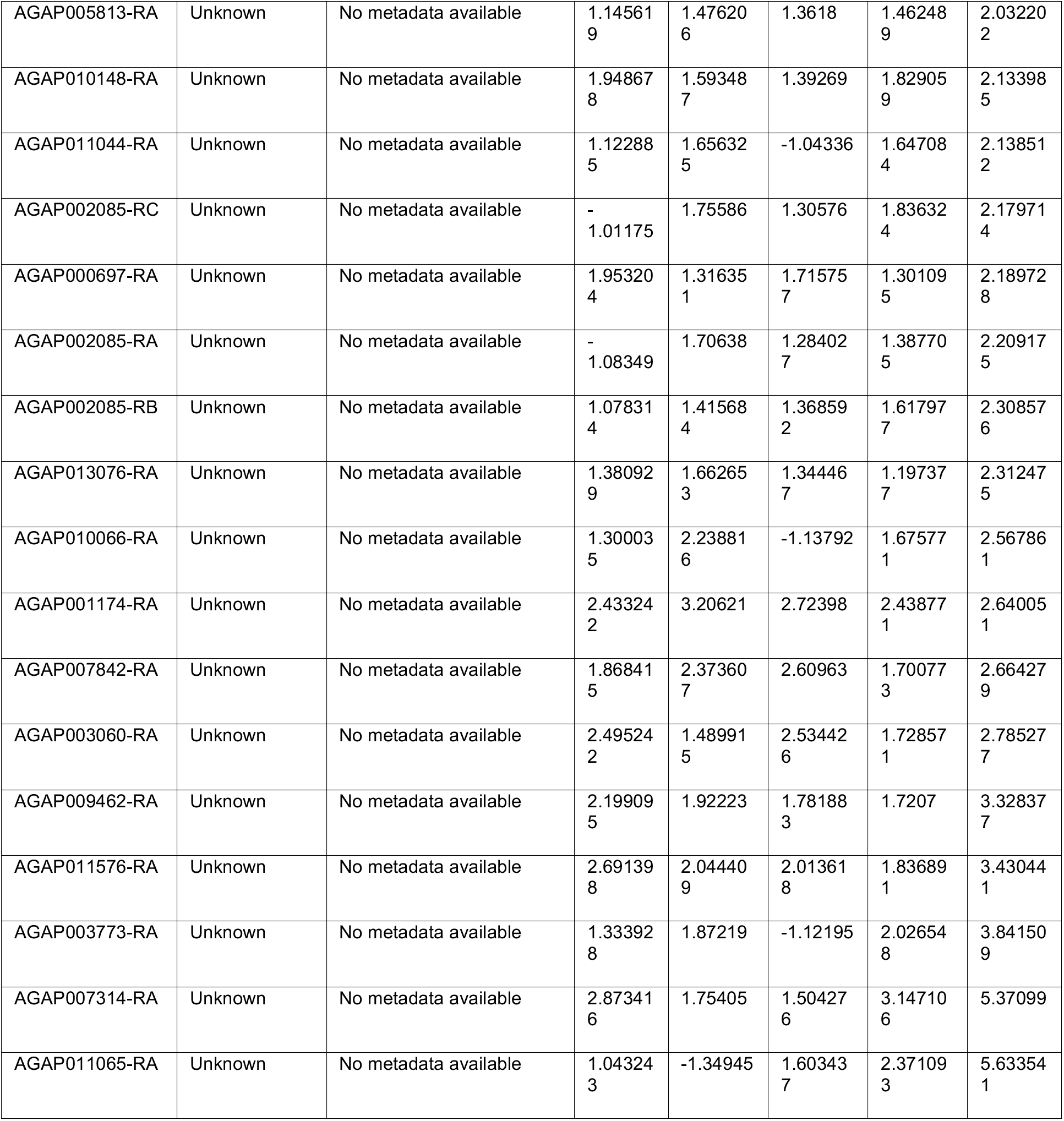
List of genes showing altered expression across time points in SCD1 KD mosquitoes.

**Table S2.**
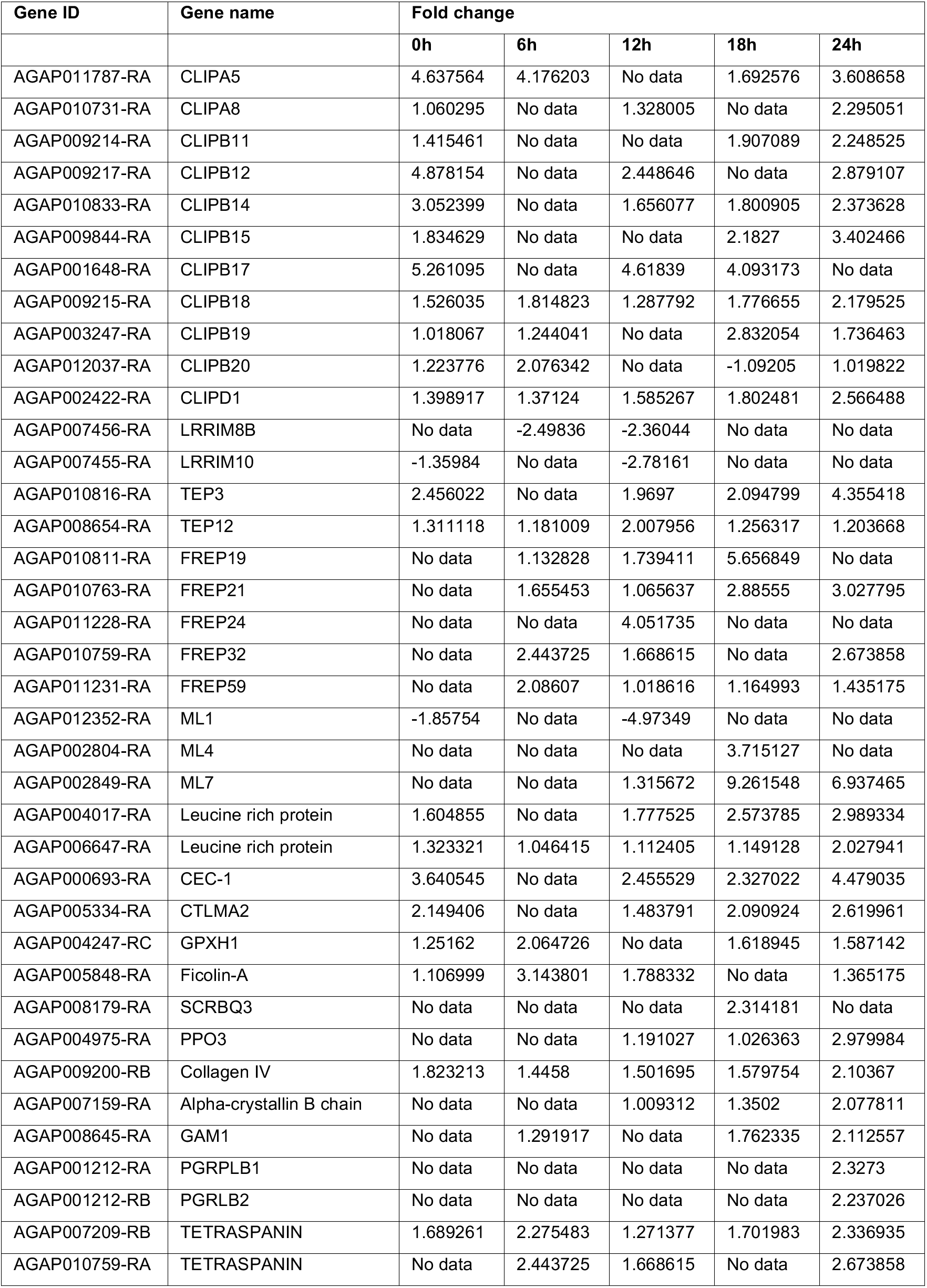
List of immune genes which show differential expression in *SCD1* KD mosquitoes.

## Notes

### Competing Interest Statement

The authors have declared no competing interest.

